# Simultaneous pathway engagement of translation in astrocytes, circadian rhythm in GABAergic neurons and cytoskeleton in glutamatergic neurons precede electroencephalographic changes in neurodegenerative prion disease

**DOI:** 10.1101/2021.11.26.470101

**Authors:** Lech Kaczmarczyk, Melvin Schleif, Lars Dittrich, Rhiannan Williams, Maruša Koderman, Vikas Bansal, Ashish Rajput, Theresa Schulte, Maria Jonson, Clemens Krost, Fabio Testaquadra, Stefan Bonn, Walker S. Jackson

**Affiliations:** Wallenberg Center for Molecular Medicine, Department of Biomedical and Clinical Sciences, Linköping University, Linköping, Sweden; German Center for Neurodegenerative Diseases, Bonn, Germany; Institute of Neurogenomics, Helmholtz Zentrum München, Neuherberg, Germany; Institute of Medical Systems Biology, Center for Molecular Neurobiology, University Medical Center Hamburg-Eppendorf; Maximon AG, 6300 Zug, Switzerland; German Center for Neurodegenerative Diseases, Tübingen, Germany

**Keywords:** translatome, gene expression, RiboTag, proteostatic stress, mitochondria, ribosome, circadian rhythm, EEG

## Abstract

Selective vulnerability is an enigmatic feature of neurodegenerative diseases (NDs), whereby a widely expressed protein causes lesions in specific brain regions and cell types. This selectivity may arise from cells possessing varying capacities to regain proteostasis when stressed by cytotoxic protein conformers. Using the RiboTag method in mice, translational responses of five neural subtypes to acquired prion disease (PrD) were measured. Pre-onset and disease onset timepoints were chosen based on longitudinal electroencephalography (EEG) that revealed a gradual increase in theta power between 10- and 18-weeks after prion injection, resembling a clinical feature of human PrD. At disease onset, marked by significantly increased theta power and histopathological lesions, mice had pronounced translatome changes in all five cell types despite having a normal outward appearance. Remarkably, at a pre-onset stage, prior to EEG and neuropathological changes, we found that 1) translatomes of astrocytes indicated a sharply reduced synthesis of ribosomal and mitochondrial components, 2) excitatory neurons showed increased expression of cytoskeletal genes, and 3) inhibitory neurons revealed reduced expression of circadian rhythm network genes. Further assessment for the role of circadian rhythms using a jet lag paradigm modestly exacerbated disease. These data demonstrate that early translatome responses to neurodegeneration emerge prior to other signs of disease and are unique to different cell types. Therapeutic strategies may need to target multiple pathways, each in specific populations of cells, early in the disease process.

## Introduction

NDs typically involve the misfolding and aggregation of proteins which lead to toxicity and cellular stress. Most NDs are characterized by late onset of clinical signs and, conceivably, protective mechanisms must be proactive to counterbalance the toxic processes that eventually manifest as disease symptoms. Neurodegeneration progresses when the capacity of those protective proteostatic mechanisms is exceeded (Fu, Hardy, & Duff, 2018). NDs initially affect defined regions in the brain, a phenomenon known as selective vulnerability (Carroll, Guha, Nehrke, & Johnson, 2021; Fu et al., 2018; Guentchev, Wanschitz, Voigtlander, Flicker, & Budka, 1999; Jackson, 2014; Mattsson, Schott, Hardy, Turner, & Zetterberg, 2016). Expression of disease-causing proteins is typically ubiquitous, and often higher in regions and cell types resistant to disease (Jackson, 2014; Kaczmarczyk et al., 2021), indicating the observed regional and cellular selectivity is determined by other factors. We hypothesize that the differences in cellular microenvironments (e.g., specific compositions of the protein trafficking and quality control machineries) influence the vulnerability of individual cells, which adopt distinct responses to cope with the disease-related protein conformers (Fu et al., 2018; Jackson, 2014; Tebbenkamp & Borchelt, 2010). As brain regions are largely composed of neural networks consisting of multiple functionally interacting cell types, failure of one cell type likely triggers a domino effect that results in region-wide changes observed in human NDs (Jackson, 2014).

In this study we dissected responses of five neural cell types to proteostatic stress using the RML (Rocky Mountain Labs) mouse model of neurodegenerative PrD. In contrast to other NDs, PrDs naturally affect several mammalian species, and mouse models present neuropathological, biochemical and gene expression changes very similar to humans. Moreover, knowledge gained by studying prion diseases is important for other NDs since the prion protein (PrP) that causes PrDs, has a mechanistic role in them (Corbett et al., 2020) and many are thought to spread through the brain via prion-like mechanisms (Goedert, Clavaguera, & Tolnay, 2010; Lau et al., 2020; Walker, Schelle, & Jucker, 2016). RML is an ideal model since it is extraordinarily precise, causing lethality to an entire study group within a week following a five-month disease course. Such precision in disease progression is well suited for gene expression analyses. We thus studied cell type-specific responses to RML using RiboTag profiling (Sanz et al., 2009). RiboTag is a Cre-directed transgenic tool that enables the isolation of ribosome-associated mRNA from cells expressing an epitope-tagged ribosomal protein (Fig 1A) (Sanz et al., 2009). By capturing actively translated mRNAs rather than total RNA, RiboTag data more closely reflect the changes in the cellular proteome of specific cell types (Battle et al., 2015). Moreover, tissue samples can be immediately frozen upon removal, constraining batch effects and preserving mRNA significantly better than methods based on physical separation of cells or cell bodies (Kang et al., 2018; Millard et al., 2021). To determine the most informative analytical timepoints, we tracked theta frequency waves, which increase in human PrD, using longitudinal EEG analysis (Franko et al., 2016). Changes to gene expression were analyzed for five brain cell types at the disease onset stage, when theta was significantly increased, and at an earlier, pre-onset stage when theta for both groups was still identical and neuropathological changes were absent.

**Figure 1.**
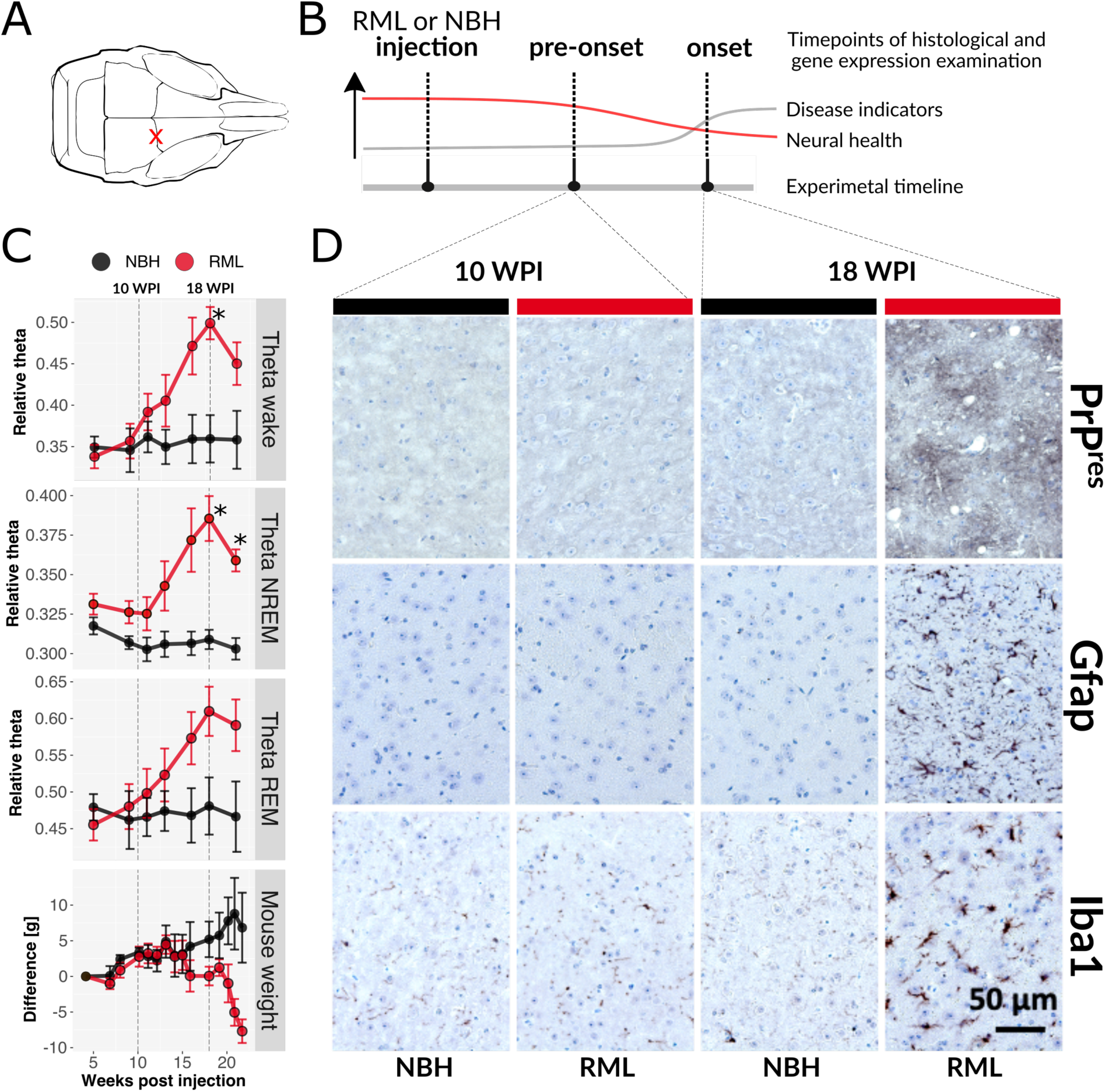
Gradual increase in EEG theta power and histopathological changes in RML. (**A**) Location of RML and NBH injections, marked by a red X. (**B**) Conceptual experimental timeline for collection of tissues for translatome and histopathological analysis, in relation to disease progression. At the pre-onset stage mice appear normal and lack disease indicators, but neural health begins to decline. (**C**) Summarized EEG theta power of wake (top), NREM sleep (second from top) and REM sleep (third from top) mice injected with RML (red) or NBH (black) juxtaposed with weekly percentage body weight measurement (expressed as deviation from initial weight) of the same mouse cohorts (bottom). Two-factor mixed ANOVA tests demonstrate significant difference in all four measurements (respectively: F(6,48) = 8.9, p-value < 0.001; F(6,48) = 6.9, p-value < 0.001; F(6,48) = 8.3, p-value < 0.001; F(14,98) = 4.8, p-value < 0.001). Differences were significant (p-value ≤ 0.05, Holm-Šídák test) at 18 WPI for wake and NREM and at 21 WPI for NREM (marked by *). (**D**) Immunohistochemistry staining of thalamus of RML infected and control (mice injected with NBH) 4 μm thick formalin fixed paraffin embedded brain sections at 10 and 18 WPI. Staining with SAF84 prion protein antibody (brown) after proteinase K treatment to detect aggregated PrP (PrP^res^, top row), staining with GFAP antibody (dark brown) to detect astrogliosis (middle row) and staining with Iba1 antibody (dark brown) to detect microglia activation (bottom row). All sections were counterstained with hematoxylin. Spongiosis is apparent in the context of the intense SAF84 staining. Scale bar represents 50 µm and applies to all images.

Even at the pre-onset stage, three cell types demonstrated coordinated, but distinct, responses although the cell type we most expected to respond did not. However, all five cell types were strongly altered at the later, disease onset stage, which coincided with increased EEG theta power. Some of the differentially expressed genes (DEGs) from the early time point have been related to PrDs and other NDs; here we present such alterations in the context of specific cell types. More significantly, this work indicates that disparate cell types unleash unique strategies in response to neurodegeneration and that parallel engagement of multiple therapeutic targets may provide the most efficacious treatments.

## Results

### A steady increase in EEG theta power marks RML progression

We have observed that a subset of mice in the commonly used C57BL/6 background are hyperactive at night (Jackson et al., 2009; Jackson et al., 2013; Kaczmarczyk et al., 2021). In contrast, 129S4 mice are calmer, exhibit more consistent behavior between individuals, and have a very different sleep pattern than C57BL/6 mice (Dittrich, Petese, & Jackson, 2017; Kaczmarczyk et al., 2021). Therefore, to optimize the identification of markers of PrD through gene expression analysis, which could be obscured by bouts of hyperactivity, we used the 129S4 mouse strain for these studies.

The neurodegenerative PrD model was initiated by unilateral, intracranial injection of brain homogenate from either normal (NBH, control) or RML infected mice (Fig 1A). RML induced terminal disease in 129S4 mice following an incubation period of 22.6 weeks, with behavioral features typical for prion diseases (e.g., kyphosis, ataxia, reduced body condition, poor grooming, clasping during tail suspension, etc.) To identify early cellular mechanisms altered by RML, we defined study stages as 1) pre-onset, before any differences and 2) onset, when differences between NBH and RML groups could be detected using a clinically relevant method in living mice (Fig 1B). This was accomplished by measuring EEG changes during sleep-wake cycles, concomitant with core body temperature (Tb) and locomotor activity in freely-moving mice implanted with wireless, telemetric devices (Data Sciences Inc). 24 h recordings were taken every two to four weeks until terminal disease was reached. Body weight was measured weekly.

EEG measures the summarized electrical activity of cortical and subcortical neurons, providing a very sensitive tool to detect brain dysfunction (Leuchter et al., 2017; Meghdadi et al., 2021), including theta power increases in several human PrDs (Franko et al., 2016). Analyses of EEG spectral frequency distributions revealed a pronounced and progressive increase in the theta band (5-10 Hz) relative to other frequencies (0-50 Hz) that reached statistical significance at 18 weeks post-injection (WPI) (Fig 1C). Theta power increase was apparent in wake, non-rapid eye movement (NREM) sleep, and rapid eye movement (REM) sleep states (Fig S1). To examine if EEG theta reflected disease progression, we calculated relative theta power (5-10 Hz/0-50 Hz) against WPI. We found that relative theta power remained consistent in controls yet incrementally increased in RML with disease progression, across all vigilance states (Fig 1C). Mixed model ANOVAs revealed interaction effects of treatment group (RML and NBH) and timepoint (WPI): wake (F_6,48_ = 8.3, p = 0.003), NREM (F_6,48_ = 6.9, p = 0.011), and REM (F_6,48_ = 8.9, p = 0.002). At 18 WPI, a peak effect was seen, reflected by statistically significant higher theta in RML mice compared to NBH mice (Fig 1C). By focusing on the evolution of EEG theta across WPI, we determined that theta power was comparable between RML and NBH mice up to 11 WPI, after which the groups slowly separated (Fig 1C).

Next, we examined brains of RML and NBH injected mice for neuropathological changes. PrDs lead to PrP aggregation (PrP^res^) and morphological changes of astrocytes and microglia readily detectable with specific antibodies (PrP, GFAP and Iba1, respectively). Each of these neuropathological hallmarks of PrD were detected in RML mice at 18 but not 10 WPI (Fig 1D). Accordingly, and combined with the EEG theta marker, 18 WPI for RML mice appeared to be a clinically relevant timepoint indicating disease onset as reflected by evident change in brain activity and histology. Nevertheless, mice demonstrated an overtly normal phenotypic behavior, with no differences found in overall sleep parameters, core body temperature, locomotor activity, or response to 6 h of sleep deprivation by the gentle handling method (Fig S1), and still lacked the typical signs of RML (e.g., kyphosis, ataxia, etc.) Lastly, by combining histological and EEG analyses, we identified 10 WPI as an early pre-onset disease time point. At this timepoint, there was no divergence in theta power or body weight (Fig 1C), and no overt histological changes (Fig. 1D) between RML and NBH mice.

### Cell types demonstrate specific responses to RML

We directed the expression of epitope-tagged ribosomes (Sanz et al., 2009) to astrocytes and four neuronal types (glutamatergic, GABAergic, PV and SST neurons) using knock-in mouse lines expressing Cre from endogenous *Gja1* (Cx43), *Slc17a6* (vGluT2), *Gad2*, *Pvalb* (PV), and *Sst* genes, respectively (Fig 2A, 2B). These cell types were chosen primarily as GABAergic neurons, particularly PV-expressing cells, are reported to be the most vulnerable in human and rodent PrDs (Ferrer, Casas, & Rivera, 1993; Guentchev, Groschup, Kordek, Liberski, & Budka, 1998; Guentchev, Hainfellner, Trabattoni, & Budka, 1997; Guentchev et al., 1999). SST neurons, while generally non-overlapping with PV neurons (Taniguchi, 2014), constitute an important GABAergic subtype. In contrast, glutamatergic neurons (excitatory) drive neural activity and the overall network excitability is balanced by GABAergic (inhibitory) populations. Lastly, we chose to study astrocytes as they play a key role in the removal of toxic substances from extracellular space (Iliff et al., 2012) and undergo reactive gliosis in PrDs and other NDs (Makarava, Chang, Kushwaha, & Baskakov, 2019; Makarava, Mychko, Chang, Molesworth, & Baskakov, 2021; Sofroniew & Vinters, 2010). An inducible Cre line was employed for astrocytes since constitutive Cre expressing lines are active in neural precursor cells and thus also label neurons (Pfrieger & Slezak, 2012).

**Figure 2.**
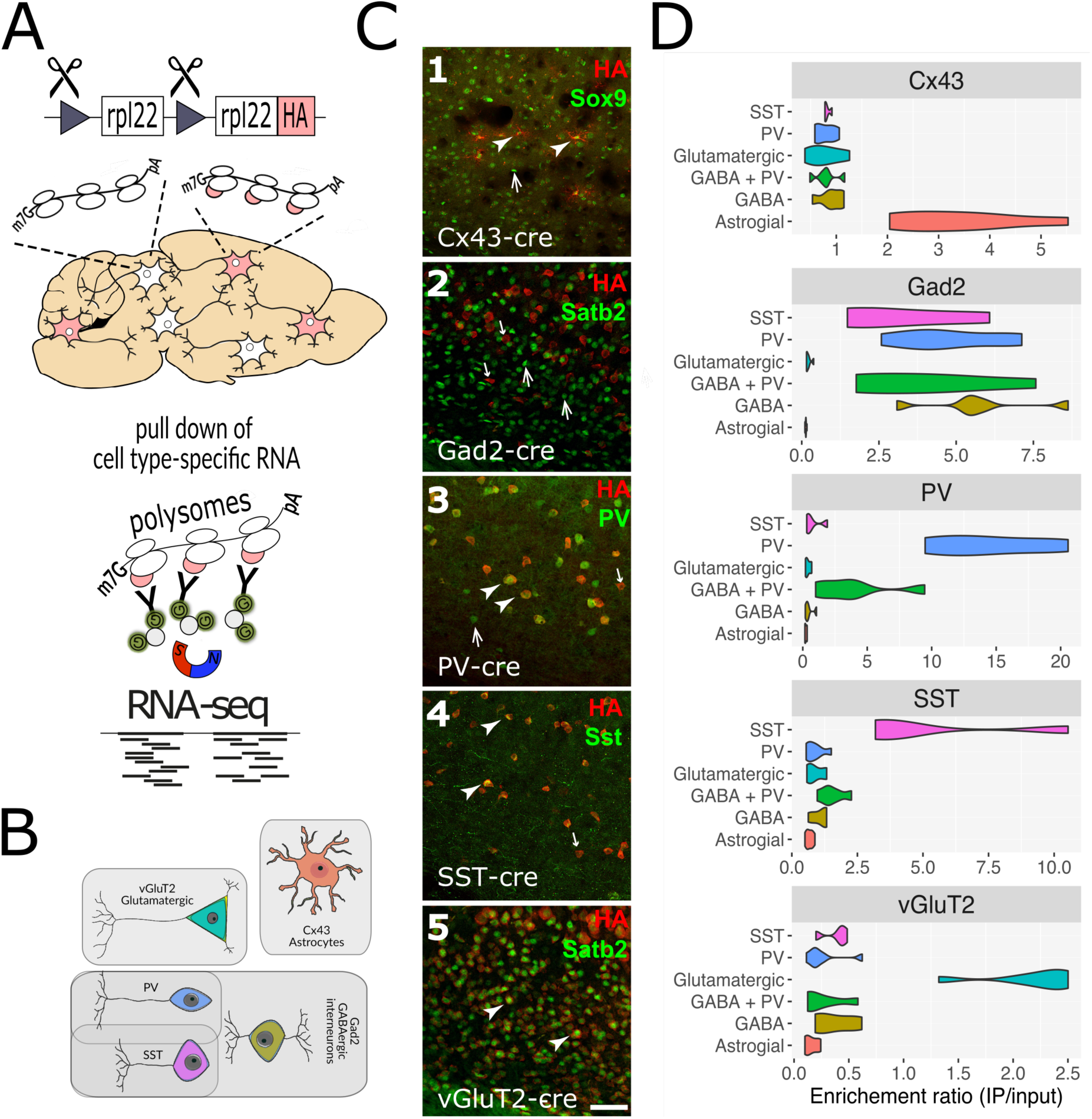
RiboTag activation and validation. (**A**) The RiboTag system is activated in cells with Cre activity through site-directed recombination in Rpl22 to make Rpl22- HA (RiboTag). Frozen mouse brain tissue is homogenized, creating a mix of mRNAs attached to either normal ribosomes (white oval bubbles) or RiboTagged ribosomes (white oval bubbles with pink tags) made specifically by Cre+ cells. Cell-type-specific, ribosome associated mRNA is isolated by RiboTag IP with HA antibody and protein G magnetic beads. mRNA from Cre+ cells is then sequenced. (**B**) Cell types used in these studies. Shaded areas reflect the expression overlap between Gad2, PV and SST Cre lines. (**C**) Confocal images of cortex following double immunofluorescence labeling of HA (RiboTag) and representative histological cell-type markers. Cell type markers used (from 1 to 5 respectively): Astrocytes (Sox9), GABAergic cells (Satb2; glutamatergic cell marker to highlight the lack of colocalization with GABAergic neurons), PV neurons (PV), SST neurons (Sst), glutamatergic neurons (Satb2). Examples for double-positive cells are indicated by arrowheads, HA-/marker+ cells by upward pointing arrows, HA+/marker- cells by downward pointing and smaller arrows. Scale bar indicates 50 µm (identical for all panels). C1: specific HA-labeled astrocyte- like shaped cells colocalize with Sox9 in Cnx43-Cre/RiboTag mice. C2: Complete separation of HA and the glutamatergic marker, Satb2, in Gad2-Cre/RiboTag mice was observed. C3: PV and HA show substantial colocalization in PV-Cre/RiboTag mice. C4: All Sst+ cells were co-labeled for HA, although there were several HA+/SST- cells in SST-Cre/RiboTag mice. C5: Nearly all cells were co-labeled for HA and the nuclear marker for glutamatergic neurons, Satb2, in the vGlut2-cre/RiboTag mice. (**D**) Violin plots showing RiboTag IP specificity; each violin summarizes absolute expression (TPM) ratios (X-axis) of known cell type markers in IP samples and corresponding input (total RNA) samples (Y-axis).

We verified the fidelity of RiboTag expression through co-localization of the RiboTag epitope with known cell type markers in histological sections (Fig 2C and Fig S2). RiboTag IP samples and corresponding total mRNA (input control) analyzed by RNA-sequencing (RNA-seq) confirmed cell type-specificity with enrichment or depletion of known cell type marker genes (Fig 2D). Principle component analysis revealed that samples from the same cell types clustered tightly together (Fig S3).

Since we expected changes to be very mild at 10 WPI, we defined differentially expressed genes (DEGs) to have a Benjamini-Hochberg adjusted p-value (hereafter false discovery rate or FDR) < 0.1 without applying a fold change cut-off. A summary of the DEGs that are unique for each cell type or shared between multiple cells, at each time point, is presented in Fig 3A (an interactive browser of all DEGs for each cell type is available online (https://shiny.it.liu.se/shiny/Scrapie_RML_RiboTag). Although at 18 WPI RML mice appeared normal via passive observation, there were severe gene expression changes for all cell types, with 8 to 40% of all mRNAs altered (FDR ≤ 0.1), and thousands of DEGs having an absolute log_2_ fold change |L2FC| ≥ 1 (Fig 3B), consistent with gross EEG and IHC (Fig 1). In contrast, at 10 WPI, SST and PV neurons in RML mice were barely altered (1 and 3 DEGs, respectively, considered unaffected cell types hereafter), contrarily to our predictions based on literature (Ferrer et al., 1993; Guentchev et al., 1998; Guentchev et al., 1997; Guentchev et al., 1999), but consistent with a recent report (Scheckel, Imeri, Schwarz, & Aguzzi, 2020). Both glutamatergic and GABAergic neurons showed more changes, with 38 and 83 DEGs respectively, but astrocytes demonstrated the greatest number of DEGs with 139 (Fig. 3A and B). Glutamatergic DEGs were primarily upregulated, whereas GABAergic and astrocytic DEGs were primarily downregulated (Fig 3A and B). Importantly, in contrast to 18 WPI when over half of DEGs were changed in multiple cell types, only 10% of 10 WPI DEGs were changed in multiple cells, indicating that early in disease, cells activate unique responses (Fig 3A).

**Figure 3.**
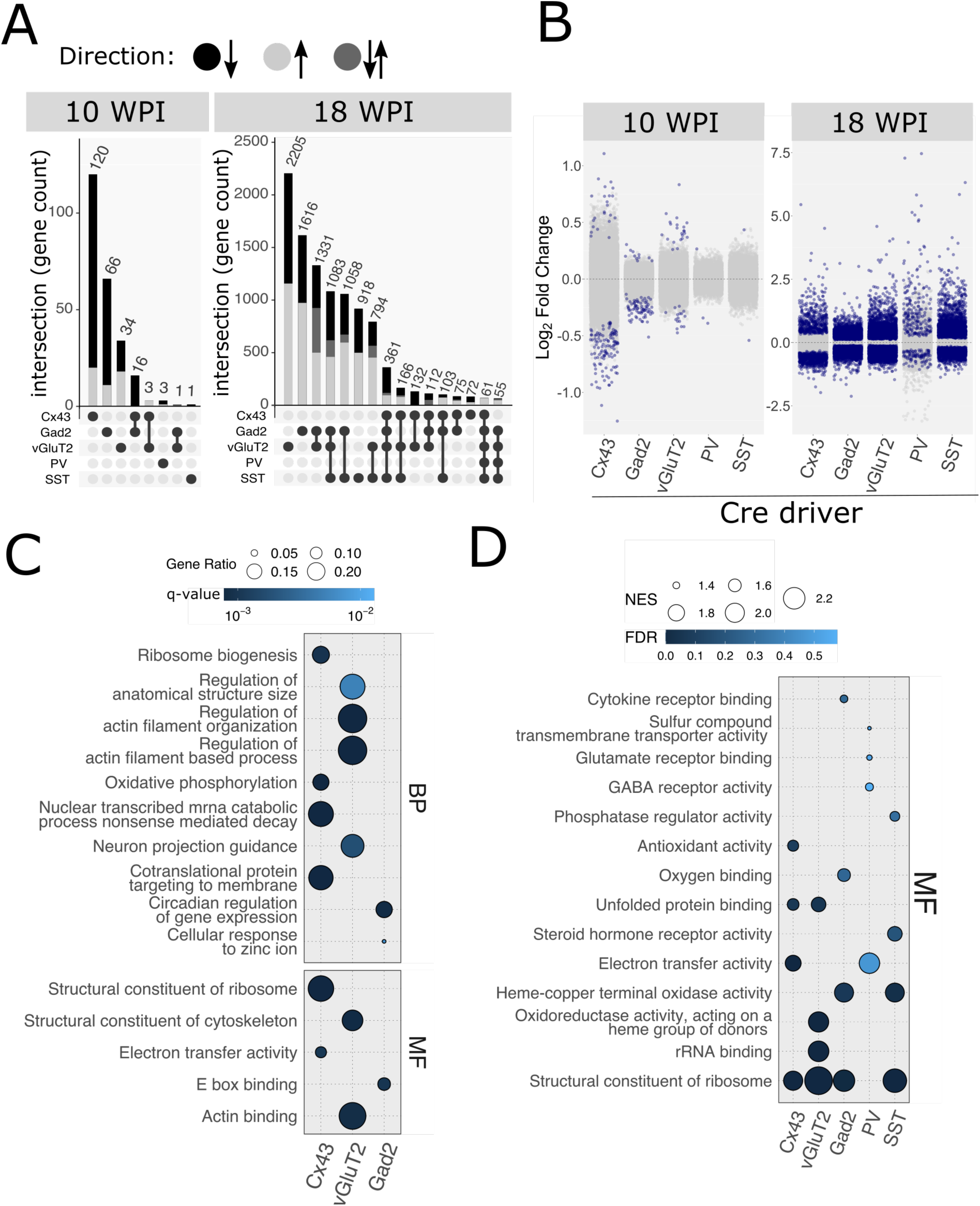
Overview of translatome changes. (**A**) Numbers of differentially expressed genes (DEGs). Values represent numbers of genes found to be differentially expressed (FDR ≤ 0.1) in a single or combination of cell types studied. Combinations are marked by dumbbell-shaped symbols below the plots. The shade of grey corresponds to the directionality of changes (see legend above the panel). For 18 WPI all clusters containing < 100 genes were omitted. Note that for 10 WPI, the changes in the 3 affected cell types: astrocytes, glutamatergic cells and GABA cells are mostly non-overlapping (respectively, 120, 66, and 34) (**B**) Visualization of DEGs (FDR ≤ 0.1; dark blue) and non-DEGs (FDR > 0.1; light grey) in all cell types and timepoints. Y-axis represents log_2_ fold change, X-axis represents cell type. (**C**) Overrepresentation analysis of DEGs in the affected cell types at 10 WPI. Significantly enriched categories were shown for molecular function (MF) and biological process (BP) gene ontology subsets. Categories related to translation, cytoskeletal regulation and circadian rhythm were the most enriched in astrocytes, glutamatergic neurons and GABAergic neurons, respectively. (**D**) Gene set enrichment analysis (GSEA) in all cell types analyzed at 18 WPI. Significantly enriched categories were shown for the MF gene ontology subset. The most prominent categories were related to translation and terminal oxidation (mitochondrial respiration). Cell type legend for all panels: Cx43: astrocytes, Gad2: GABAergic cells, vGluT2: glutamatergic neurons, PV, parvalbumin neurons, SST: Somatostatin neurons.

We expected that if cell responses were deliberate, multiple DEGs for a given cell type to be associated with a specific cellular pathway or process. Since the data from the two timepoints have vastly different numbers and magnitudes of DEGs, we optimized our analysis by using the most appropriate method for the properties of the data. Since the 18 WPI data were characterized by a large percentage of genes being differentially expressed, and consequently multiple functional modalities in the cells were affected, we analyzed those data with gene set enrichment analysis (GSEA). GSEA considers the ranking of all genes (DEGs and non-DEGs), thereby maximizing the potential to determine the most affected modalities (higher gene ranking contributes more to the overall GSEA scoring). In contrast, overrepresentation analysis (ORA) uses unranked DEG subsets which is more appropriate for the 10 WPI data since relatively small changes may not be appropriately reflected in the ranking due to a lower signal-to-noise ratio. Genes were ranked based on the degree of differential expression and log_2_fold change (L_2_FC) using the formula: *`rnk`* = *LFC*\*(−log(*FDR*)). Noteworthy, when applied to 10 WPI sample data, GSEA failed to yield significant enrichment scores. Corroborating the differences between the cell types with respect to differential gene expression (Fig 3A), this analysis further confirmed that there were no common pathways at 10 WPI (Fig 3C), although the cells responded similarly at 18 WPI, with ribosomal and electron transfer molecular functions in common (Fig 3D, S4). Intrigued by the changes at 10 WPI, we focused the remainder of our analysis on those data. To explore specific cellular changes further, we included interactome information in our analysis.

### DEGs occupy neighboring positions in the interactome

By identifying genes in regions of the interactome connected to DEGs, we could deduce which cellular changes are relevant to disease progression. The proteins encoded by genes sharing disease associations often colocalize within protein-protein interactions (PPI) networks, forming communities, referred to as disease modules. Identification of disease modules typically relies on exploring the interactions of known disease genes, i.e., the seed genes. Herein, we defined the seeds as the subset of gene network nodes overlapping with DEGs and examined whether they provide sufficient clues on disease module localization. To this end we collected high confidence PPI interactions (cumulative confidence score > 0.7) from the STRING database and constructed a separate interactome for each cell type, excluding the genes that were not detected in either diseased or control mice. We then determined the sizes of the largest connected components (LCCs) within subnetworks composed of seed genes and compared each to a reference LCC size distribution of randomly selected nodes in the interactome. In astrocytes, GABAergic, and glutamatergic cells the LCC sizes were greater than for random subnetworks of equivalent size (Z-scores 2.53, 5.42, 2.57 respectively, empirical p-values 0.005, 0.001, 0.043) (Fig S5A). The general lack of DEGs in PV and SST neurons was corroborated with a lack of significant connectivity between the top ranked genes (0.01 Wald test p-value cutoff was used) in those cell types (Z-scores 0.30 and 1.69 respectively, empirical p-values 0.342 and 0.077, Fig S5B).

We then sought to identify putative disease modules specific to each affected cell type in early pre-onset disease stage (10 WPI) by integrating the interactome and RiboTag data sets using a modified DIAMOnD method (Ghiassian, Menche, & Barabasi, 2015). To account for bias towards well studied genes in the STRING-based network, our modified version of the DIAMOnD algorithm used a two-stage ranking system to iteratively add genes to a disease module (see supplementary materials and methods). Identified disease modules for all cell types were enriched with genes which share functional annotations with seed genes and therefore represent network communities likely affected by disease. The results are described in the following sections, organized by cell type.

### Astrocytes downregulate ribosomal and mitochondrial mRNAs

The most prominent feature of DEGs in astrocytes was a strong down regulation of mRNAs encoding 26 ribosomal proteins, (average L_2_FC = -0.64; range -0.41 to -0.94). Additional proteins related to ribosome synthesis (Fbl) or translation in general (Ccdc124, Denr, Dohh, Eif5b, Upf2) were also downregulated, supporting the view that fewer ribosomes were being made. Mitochondrial proteins (Glrx5, Lars2, mRpl18, mRps12, Romo1, Tomm6) and components of the electron transport chain including cytochrome c, and proteins in complexes I (n=3), III (n=4), IV (n=3) and V (n=3), were all downregulated in astrocytes, except for Lars2 which was upregulated. The most strongly downregulated gene, Pcsk1n (L_2_FC -1.25, FDR < 3E-5), encodes proSAAS, a secreted chaperone that has anti-aggregation activity and is found associated with protein aggregates in brains of Alzheimer’s disease (AD) patients and an AD mouse model (Hoshino et al., 2014). Interestingly, RML induced a decrease in macrophage migration inhibitory factor (Mif), within astrocytes. This factor is induced in neurons in AD patients and mouse models (Zhang et al., 2019).

There were also three up-regulated DEGs of interest: C4b, Serpina3n, and Tnrc6a. C4b is a marker for aging astrocytes and signals for inflammation (Boisvert, Erikson, Shokhirev, & Allen, 2018), Serpina3n is also a marker for aging astrocytes (Boisvert et al., 2018) and is strongly up-regulated in all forms of human PrDs (Vanni et al., 2017), and Tnrc6a associates with a complex involving PrP and Argonaute2 for a role in miRNA-mediated silencing (Gibbings et al., 2012).

These observations of 10 WPI DEGs being closely related to ribosome biogenesis and oxidative phosphorylation were confirmed by our ORA analysis (Fig 4A). IHC experiments verified Rps21 was reduced by RML at 10 WPI in cerebellar astrocytes (Fig 4B) which preferentially express Rps21 (Kaczmarczyk et al., 2021). Notably, numerous components of ribosomes and mitochondria were reduced in postmortem human PrD brains (Ansoleaga et al., 2016).

**Figure 4.**
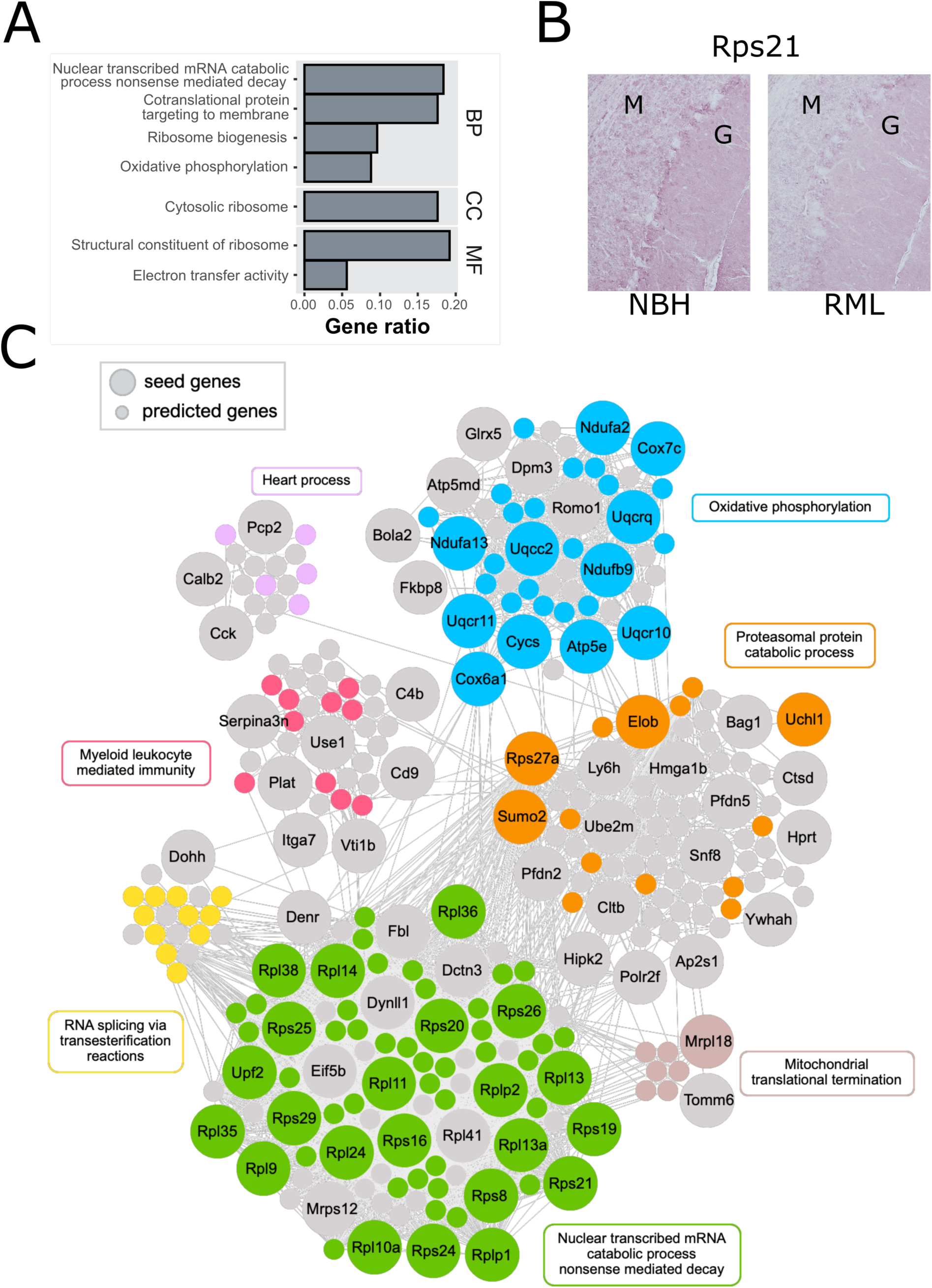
Changes in astrocytes (RiboTag::Cx43-Cre). (**A**) Overrepresentation analysis of DEGs in astrocytes (Cx43+ cells) at 10 WPI. Significantly enriched GO categories were shown for molecular function (MF), biological process (BP) and cellular component (CC) GO subsets. Most significantly enriched categories were related to translation (e.g., ribosome biogenesis) and terminal oxidation (e.g., electron transfer). X-axis is the reciprocal of gene ratio. (**B**) RML causes reduced staining of Rps21, one of the ribosomal proteins downregulated at 10 WPI, especially in Bergmann glia between molecular (M) and granular (G) cell layers. (**C**) Visualization of 10 WPI disease DIAMOnD modules constructed around 64 of 128 DEGs (large circles) in astrocytes. Small circles denote genes added by the DIAMOnD algorithm (see also Fig S6 for the same visualization with gene labels). The module consists of 6 functionally related clusters (identified with the fast greedy modularity optimization algorithm). In each cluster, the genes annotated to the representative GO term are colored.

128 DEGs from astrocytes were included in our astrocyte cell interactome and used as seed genes for inference of disease-relevant genes using the modified DIAMOnD, producing a module with 7 major communities. Within the two largest communities, the most overrepresented functional categories were ‘oxidative phosphorylation’ and ‘nuclear transcribed catabolic process nonsense mediated decay’, which consisted primarily of mitochondrial and ribosomal protein gens, respectively. In small communities, the overrepresented categories were ‘proteasomal protein catabolic process’, ‘RNA splicing via transesterification reactions’, ‘heart process’, ‘myeloid leukocyte mediated immunity’ and ‘mitochondrial translational termination’ (Fig 4C). Thus, at 10 WPI, astrocytes primarily downregulated genes for mitochondrial and ribosomal biogenesis in response to RML.

### Glutamatergic neurons upregulate cytoskeleton genes

Of the 38 DEGs altered by RML in glutamatergic neurons, many were linked to upregulation of cytoskeleton-shaping proteins (Gsn (gelsolin), Limch1, Mprip, Ppp1r9a) and cytoskeletal components including GFAP and four spectrins (Sptan1, Sptb, Sptbn1, Sptbn2) (Fig 5C). In human PrD, mass spectrometry analysis of synaptasomes revealed Gsn increased while Sptan1 decreased (Gawinecka et al., 2013). Gesolin and spectrin breakdown products are common in human NDs (Ji, Chauhan, Wegiel, Essa, & Chauhan, 2009; Yan, Jeromin, & Jeromin, 2012) and the mRNA upregulation seen here may be compensatory. The GABA receptor protein Gabrb2 was decreased which may lead to increased neuronal activity. Consistent with this notion, Arc (activity-regulated cytoskeleton) commonly induced for synaptic plasticity (Steward, Wallace, Lyford, & Worley, 1998), was increased, which in turn may have led to the increased synthesis of cytoskeleton related proteins. The ORA analysis confirmed this notion with biological process terms including ‘regulation of long-term synaptic depression’, ‘membrane raft organization’, and ‘endoplasmic reticulum to Golgi vesicle mediated transport’ (Fig 5A).

**Figure 5.**
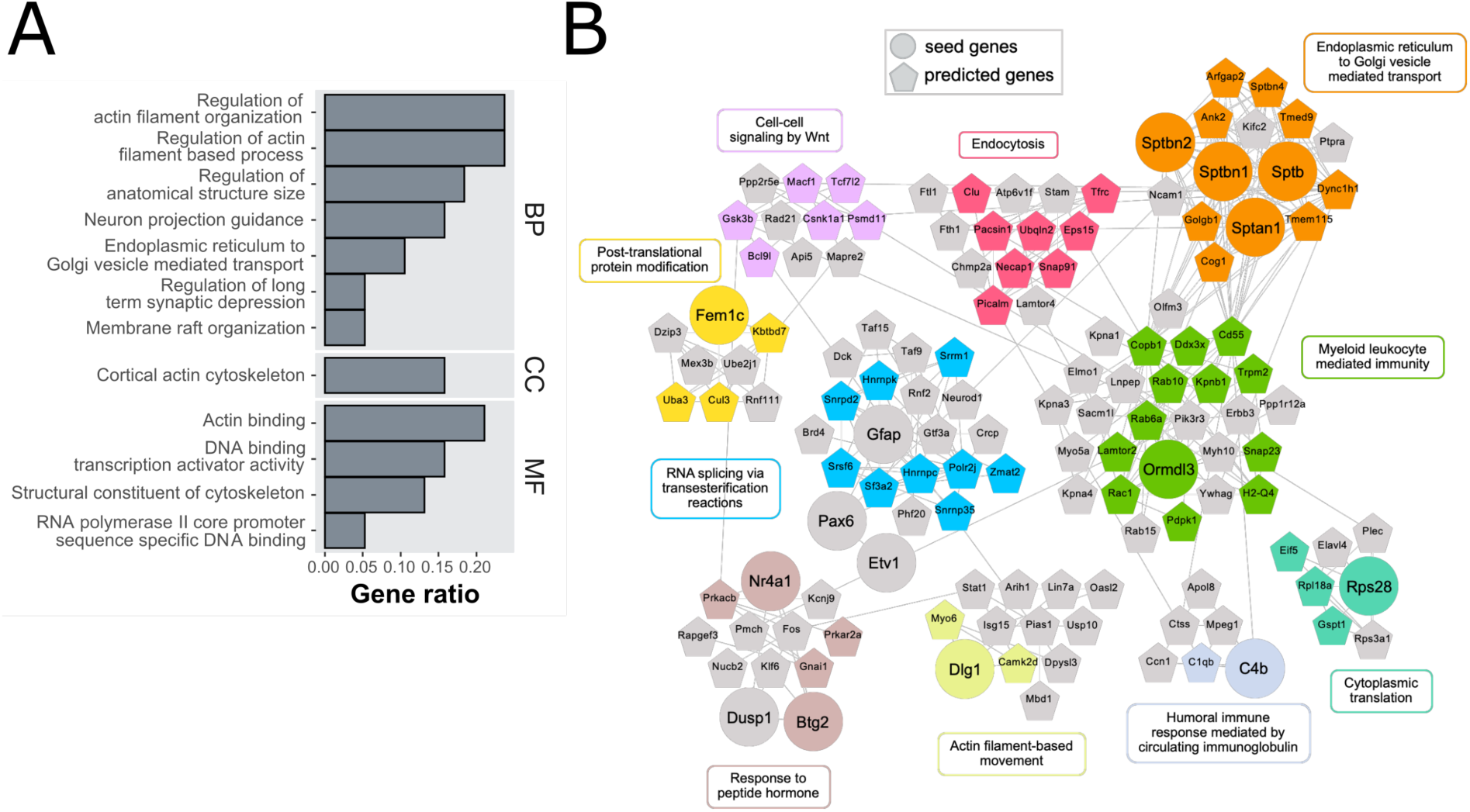
Changes in glutamatergic neurons (RiboTag::vGluT2-Cre). (**A**) Overrepresentation analysis of DEGs in glutamatergic neurons (vGluT2+ cells) at 10 WPI. Significantly enriched GO categories were shown for molecular function (MF), biological process (BP) and cellular component (CC) GO subsets. Most significantly enriched categories were related to cytoskeleton (e.g., regulation of actin filament organization) and synaptic function (e.g., regulation of long-term synaptic depression). X-axis is the reciprocal of gene ratio. (**B**) Visualization of disease DIAMOnD modules constructed around 15 of 33 DEGs (circles) in glutamatergic neurons. Pentagons denote genes predicted by the DIAMOnD algorithm. The module consists of 7 functionally related clusters (identified with the fast greedy modularity optimization algorithm). In each cluster, the genes annotated to the representative GO term are colored.

We next used the modified DIAMOnD algorithm for inference of disease-relevant genes which incorporated 33 of 38 DEGs in the glutamatergic cell interactome as seed genes. The resulting module contained 10 communities, within which the most overrepresented functional categories were ‘ER to Golgi vesicle mediated transport’, ‘RNA splicing via transesterification reactions’, ‘myeloid leukocyte mediated immunity, ‘endocytosis’, ‘response to peptide hormone’, ‘actin filament-based movement’, ‘humoral immune response mediated by circulating immunoglobulin’ ‘cytoplasmic translation’, ‘post-translational protein modification’ and cell-cell signaling by Wnt’ (Fig 5B). Therefore, at 10 WPI the changes observed in vGluT2+ cells were coordinated and quite different from those in Cx43+ cells.

### GABAergic neurons modulate circadian rhythm genes

While RML induced astrocytes to downregulate ribosomal and mitochondrial biogenesis and glutamatergic neurons increased cytoskeleton genes at 10 WPI, GABAergic neurons modulated 11 genes implicated in circadian rhythmicity: Bhlhe41 (Dec2), Ciart (chrono), Dbp, Nr1d1, Nr1d2, Per1, Per2, and Usp2, were down-regulated while, Arntl (Bmal1), Tardbp, and Vip were up-regulated. These differences are related to RML and not time of death, as the latter was controlled for in our experimental design. Remarkably, circadian rhythm genes are commonly dysregulated in neurodegenerative diseases (Carter, Justin, Gulick, & Gamsby, 2021). Many of the downregulated genes normally suppress Arntl expression, potentially explaining its upregulation. Arntl forms dimeric complexes with Clock or Npas2 to drive expression of circadian clock and redox genes, and its disruption in mice leads to neurodegeneration with prominent reactive astrocytosis (Musiek & Holtzman, 2016; Musiek et al., 2013). Studies have highlighted a role for the circadian rhythm network in antioxidation and neuroprotection (Musiek & Holtzman, 2016) and we see a potential connection in our data with down regulation of proteins involved in mitochondrial morphology (ROMO1) and electron transport complex III (Uqcc2 and Uqcrq). It is worth noting that timing may be modified by RML independently to sleep if driven by GABAergic cells.

The most significant pathways from ORA included ‘RNA catabolic process’, and ‘circadian regulation of gene expression’ for BPs, ‘postsynaptic density membrane’ for CC, and ‘structural constituent of ribosome’ for MF (Fig 6A). Analysis of 10 WPI data with the modified DIAMOnD resulted in a module built on 69 of 83 DEGs with 6 communities with functional categories of ‘ATP synthesis coupled electron transport’, ‘cotranslational protein targeting to membrane’, ‘RNA splicing’, ‘protein polyubiquitination’, ‘rhythmic process’, and ‘myeloid leukocyte mediated immunity’ (Fig 6B). Thus, although Gad2+ cells also changed expression of some ribosomal and mitochondrial proteins, they predominately changed additional pathways and demonstrated an overall response quite different from Cx43+ and vGluT2+ cells.

**Figure 6.**
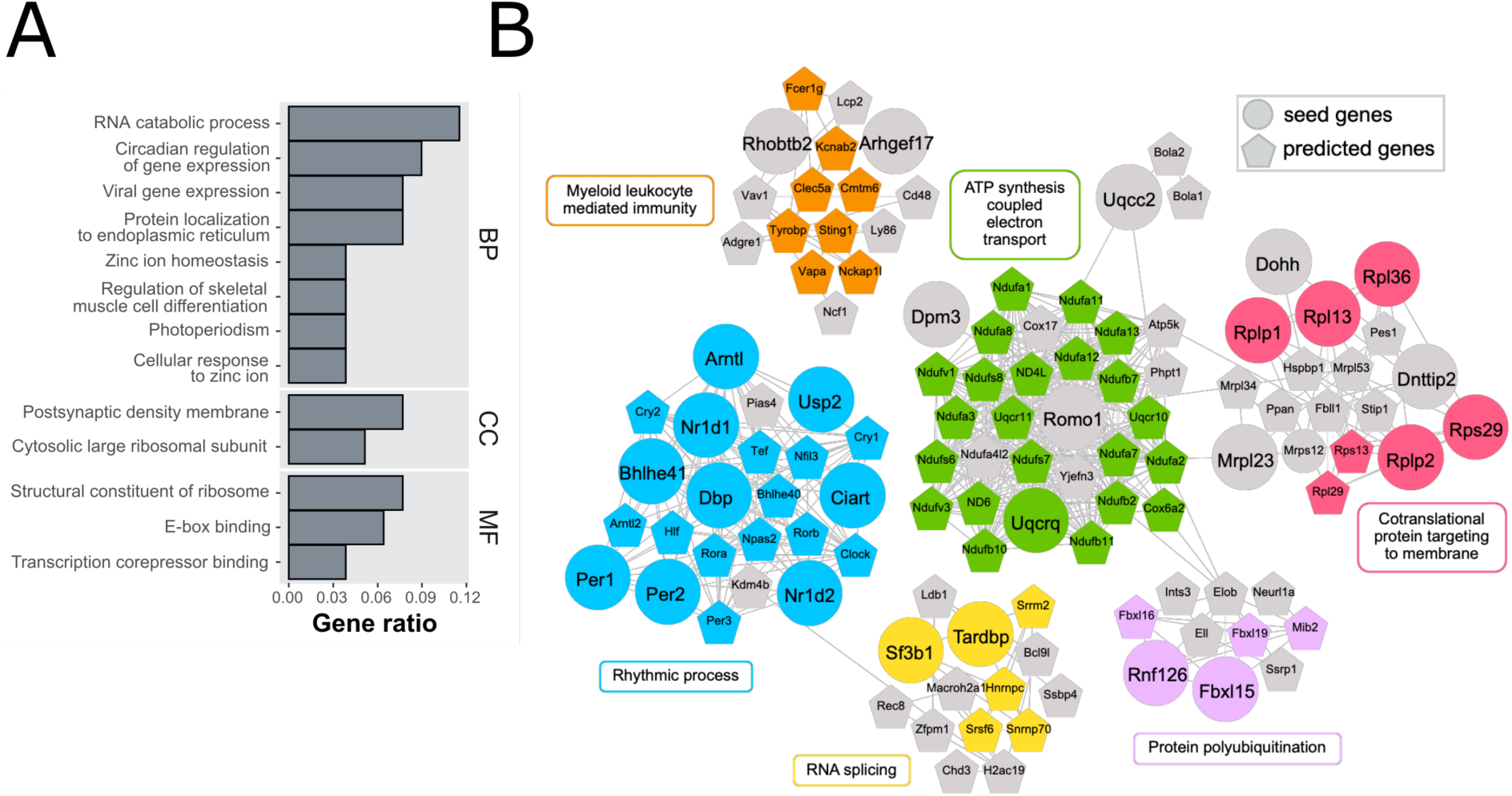
Changes in GABA neurons (RiboTag::Gad2-Cre). (**A**) Overrepresentation analysis of DEGs in GABAergic neurons (Gad2+ cells) at 10 WPI. Significantly enriched GO categories were shown for molecular function (MF), biological process (BP) and cellular component (CC) GO subsets. Most significantly enriched categories were related to circadian rhythm (e.g., circadian regulation of gene expression) and translation (e.g., cytosolic large ribosomal subunit). X-axis is the reciprocal of gene ratio. (**B**) Visualization of disease DIAMOnD modules constructed around 27 of 69 DEGs (large circles) in GABAergic neurons. Small circles denote genes added by the DIAMOnD algorithm. The module consists of 6 functionally related clusters (identified with the fast greedy modularity optimization algorithm), the largest of which involved ‘rhythmic process’. In each cluster, the genes annotated to the representative GO term are colored.

### Jet lag induced disruption of circadian rhythm mildly exacerbates RML

Since key circadian rhythm genes were changed in a key neuronal population in our study, and in several others (Carter et al., 2021), we asked if circadian rhythm regulation becomes more vulnerable to external disturbances in RML, accelerating disease progression. A common alteration of circadian rhythmicity in humans is when crossing different time zones, the poor adaptation referred to as jet lag (JL). Therefore, we subjected both RML and NBH mice to a JL modeling procedure whereby lights turn on and off 6 hours earlier each week (phase advance, PA) starting at 6 WPI. We hypothesized that if RML destabilized circadian rhythm mechanisms, this mild JL intervention might enhance RML induced theta EEG increase, or enhance reactive astrocytosis, since Arntl knockout mice demonstrate severe reactive gliosis (Musiek & Holtzman, 2016; Musiek et al., 2013).

The activity of JL mice was highest when lights were off, even as the timing shifted, demonstrating the mice sensed and responded to the PA (Fig 7A). Since comparing theta between acutely jet-lagged and undisturbed mice might be confounded by disease-independent effects of JL, we made sure there was sufficient recovery time. At 13 WPI, two weeks following the final PA, theta tended to increase, although not significantly (Fig 7B). To determine if JL enhanced reactive gliosis, two microscopists blinded to the sample identities independently examined and scored samples for severity of reactive gliosis at 14 WPI. In most comparisons, though not all, the JL samples were more intensively stained (Fig 7C). One microscopist scored the JL samples as being most severely affected in six of the seven blocks (p = 0.0101, Fisher’s exact) while the second examiner identified five (p = 0.1319, Fisher’s exact). Thus, although theta was not strongly affected, JL appeared to enhance the ongoing neurodegeneration.

**Figure 7.**
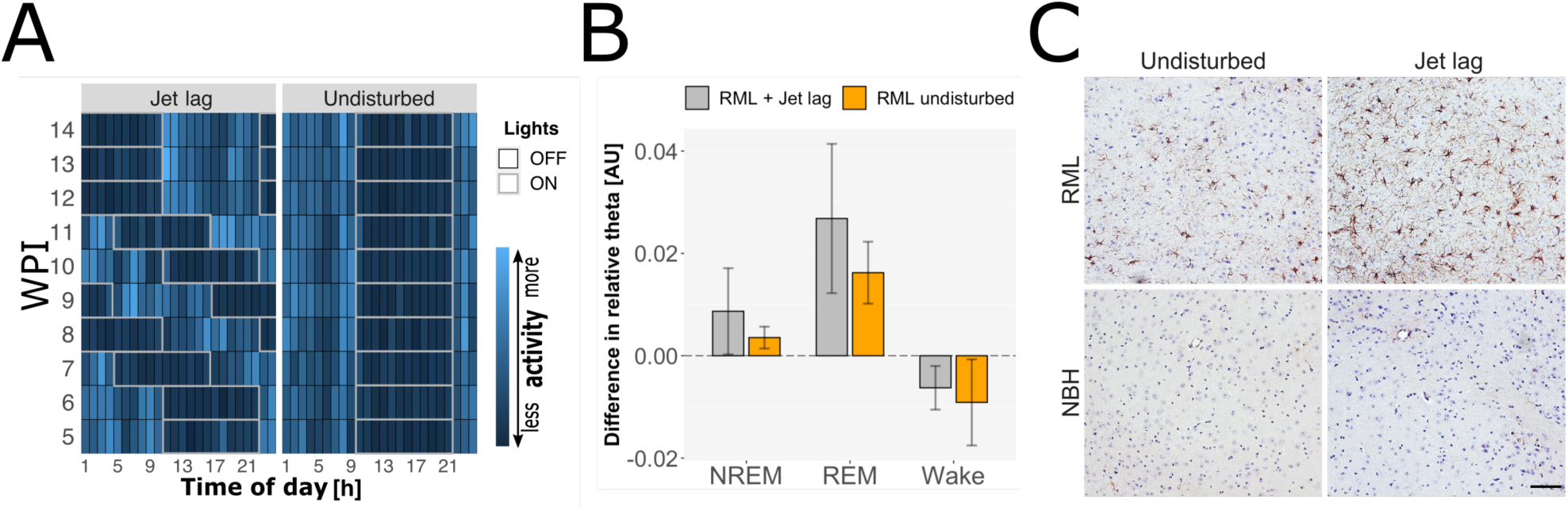
A mild jet lag challenge mildly worsens RML. (**A**) Activity (group average) of mice measured with the DSI telemetric transmitter, simultaneously recording EEG, indicates that mice are least active (dark blue tiles, each tile = 1 h) when lights are on (tile groups framed in grey) and most active (light blue tiles) when lights are off (tile groups framed in black). Each row represents data from the indicated WPI. The right panel represents the undisturbed group; note that the grey (lights on) frame for each row is aligned with the grey frames in the other rows. The left panel shows that in the Jetlag group after baseline recordings at WPI 5 and 6, the grey frames shift (advance) 6 h earlier each week until the final shift at WPI 12. Note that the tiles inside are still dark blue. (**B**) After two weeks to recover from the final PA, 24h measurements of theta at the end of WPI 13 revealed a trend consistent with enhancement of disease, although statistically not significant (ANOVA followed by Holm-Šídák test). (**C**) Thalamus stained for GFAP shows that JL modestly enhanced reactive gliosis in RML (top right versus top left) but had no effect on controls (bottom). Scalebar is 100 µm and applies to all images.

## Discussion

### Identification of EEG biomarker that precedes morphological disease changes

A better insight into mechanisms of selective vulnerability in neurodegenerative diseases is needed to develop therapies. For these therapies to stop or slow disease progression, it is important to understand disease mechanisms at early stages, before clinical signs. We chose mouse RML prion disease as a model as the timeline from induction until clinical illness and neuropathological changes is highly precise (Watts & Prusiner, 2014). To define disease onset, we measured theta waves longitudinally with EEG. As in human PrDs (Franko et al., 2016) we found theta increased. Despite spectral changes in this band, and other spectral composition alterations at 18 WPI, we did not find changes in sleep-wake states, which suggests for this RML model, that degeneration remains too mild to affect sleep or that compensation mechanisms are in place that protect sleep-wake behavior up to 18 WPI. Nonetheless, the observed increase in theta as disease progressed might be suitable for detecting effects of disease-modifying experimental treatments in mice.

### RiboTag elucidates cell type-specific responses to early ND

RML in RiboTag mice permitted the detection of early molecular responses elicited by different cell types. To verify if the RiboTag IPs contained mRNAs from the cells of interest we assessed the enrichment or depletion of known cell type marker genes compared to the total RNA input. Cell type markers are generally not absolutely specific for one cell type, but high abundance can predict the population of interest. GFAP is a notable example; as it is widely recognized as an astrocyte marker, but we found expressed in our neuronal samples, and RML induced a significant upregulation of GFAP in glutamatergic neurons at 10 WPI. Neuronal expression of GFAP has been reported by chromatin marks in healthy young adult mouse brains (Halder et al., 2016), human AD by IHC (Hol et al., 2003) and single nucleus RNAseq (Mathys et al., 2019), in motor neurons in a mouse model of amyotrophic lateral sclerosis (Sun et al., 2015), and in synaptasomes of human PrD (Gawinecka et al., 2013). Therefore, increased GFAP expression in neurons seems common to many neurodegenerative diseases. Furthermore, since several other astrocyte markers were strongly depleted/absent in neuron-derived samples, we conclude that this GFAP observation was not due to cross-contamination.

Unexpectedly, at 18 WPI there were extensive changes in gene expression even though by passive observation RML mice had no overt phenotype. This implies that even when the first clinical symptoms appear in patients of NDs, there may already be significant cellular perturbations, across several cell types making successful treatment interventions difficult. Furthermore, at 10 WPI, prior to EEG and histological abnormalities, we identified that astrocytes, glutamatergic, and GABAergic cells (presumably PV- and SST-) were responding. While at 18 WPI each cell type changed similar molecular pathways and sets of genes, at 10 WPI each population demonstrated unique DEGs and pathways and was, therefore, altered specifically. These results support our hypothesis that the phenomenon of selective vulnerability is caused by specific brain cells adopting unique responses to cope with an emerging neurodegenerative disease.

### Distinct translatome responses in excitatory and inhibitory neurons

Glutamatergic neurons at 10 WPI altered their translatomes to maintain or enhance actin and cytoskeleton networks, which are important for cellular transport to the synapse and signal transduction. Moreover, Arc responds to synaptic activity by driving expression of cytoskeletal proteins at synapses (Minatohara, Akiyoshi, & Okuno, 2015), and its upregulation suggests glutamatergic neurons are abnormally active, which may relate to excitotoxicity (Chen, Zhu, Wang, & Hang, 2020), a molecular mechanism long postulated to be involved in neurodegenerative diseases.

GABAergic neurons prominently modulated their translatomes for the circadian rhythm pathway. Since circadian rhythm genes have been associated with neurodegeneration (Carter et al., 2021; Fifel & Videnovic, 2020; Leng, Musiek, Hu, Cappuccio, & Yaffe, 2019), we tested whether interfering with the circadian rhythm system would worsen disease. Theta in RML mice measured two weeks after the last PA tended to be different between JL and undisturbed control mice but was not significant. Notably, this corresponded to 13 WPI, still before theta is increased in RML versus controls, so JL would need to have a big impact to yield a detectable difference.

In this light, we were somewhat surprised to see even the modest effect on reactive gliosis. It is conceivable that a more intense jet lag paradigm or an experiment ending later in the disease course would have a larger impact. The results presented here provide an initial framework to guide future experiments testing the effects of circadian rhythm interference on neurodegeneration.

### Astrocytes reduce synthesis of ribosomes and mitochondria

Human PrDs are characterised by reduced synthesis of mitochondria and ribosomes (Ansoleaga et al., 2016; Frau-Mendez et al., 2017), and our cell-type-specific data suggest that during early disease, these changes occur in astrocytes, even before other signs of disease. There are several possible explanations for why astrocytes deploy this strategy.

The strong reduction of ribosome protein mRNAs could be related to ribosome specialization, where the stoichiometric ratio of ribosomal proteins differs between tissues and brain regions, possibly to steer the preferential translation of specific mRNAs (Jackson, 2014; Kaczmarczyk et al., 2021; Preiss, 2016; Shi et al., 2017; Xue & Barna, 2012). However, the reduction in many ribosome proteins in combination with reduced mitochondrial proteins suggests it is not simply ribosome specialization. Since ribosomes are highly abundant and energy intensive to make, their synthesis is tightly coupled to that of mitochondria, and the reduction in both implies that complete ribosome synthesis is decreased. In the mouse brain, the half-life for ribosomal and mitochondrial proteins is approximately 10 and 20 days, respectively (Fornasiero et al., 2018). Thus, while acute intracellular signaling (e.g., phosphorylation), occurs without mRNA expression changes, drivers of long-lasting reduction in mitochondria and ribosomes will likely be directed by gene expression changes, which might be reflected in our data.

Glycogen metabolism, a key function of astrocyte energetics, was not affected as adenylate cyclases, phosphodiesterases and other genes associated with protein kinase A signaling were unchanged. The mTOR (mechanistic target of Rapamycin) pathway also senses energy levels and controls growth, in part through coordination of ribosome biogenesis. However, the only DEG related to mTOR, Fkbp8, is an inhibitor and its decrease should therefore stimulate ribosome biogenesis. Thus, a lack of glycogen and mTOR signaling changes suggest there is not a change in metabolism. The unfolded protein response acts to relieve protein misfolding stress by reducing overall protein synthesis, especially ribosomes, and increasing production of protein quality control proteins (e.g., chaperones) (Hetz, Zhang, & Kaufman, 2020). Such a response was noted in an over-expression transgenic model of acquired prion disease (Moreno et al., 2012; Smith et al., 2020). However, data from another study (Scheckel et al., 2020) and ours did not demonstrate increased synthesis of protein quality control genes, suggesting that an artifactual unfolded protein response may have been triggered in the overexpression transgene study (Smith et al., 2020).

A final explanation for the reduced ribosome synthesis in our study relates to molecular crowding. Cells tune ribosome abundance to sculpt their physical properties, including diffusion rates of intracellular protein aggregates or cellular stiffness (Delarue et al., 2018; Evers, Holt, Alberti, & Mashaghi, 2021). It is conceivable that astrocytes would benefit from such a manipulation in early stage PrD.

### These results in the context of previous studies

Previous reports have also contributed to our understanding of selective vulnerability by applying alternative methods. The laser capture microdissection approach was used to cleanly remove sections of the CA1 region of the hippocampus or granule cells of the cerebellum from mice at various stages of PrD (Majer et al., 2012; Majer et al., 2019). Purified RNA was then measured with microarrays. These regions primarily, or exclusively, contain glutamatergic neurons and are thus most comparable to our glutamatergic RiboTag samples. Interestingly, they found RML upregulated genes related to actin-cytoskeleton in both regions early in disease, with a stronger response in the cerebellum. Our data also hint at the cerebellum being the location of affected glutamatergic neurons as some of the DEGs (e.g., Zic1 and Zic2) are preferentially expressed there, according to the Allen Brain Atlas.

Another study employed fluorescent activated cell sorting to study microglia from prion diseased mice (Vincenti et al., 2015), a cell type providing an important neuroprotective function to brains during prion diseases (Zhu et al., 2016). Interestingly, microglia sharply increase expression of mitochondrial and ribosomal protein encoding genes (Vincenti et al., 2015), in direct contrast to what we report here for astrocytes. This difference may relate to the concept that ribosome abundance is employed to regulate intracellular crowding (Delarue et al., 2018; Evers et al., 2021) and/or due to differences in phagocytic capacity by micro- and astroglia during PrDs (Sinha et al., 2021).

During the completion of our work a report of experiments like ours was published (Scheckel et al., 2020). They employed a Cre dependent ribosome tagging system analogous to RiboTag to study the responses of excitatory and PV neurons, astrocytes, and microglia to intraperitoneal RML injection. Despite the overall similarity of the experiments, the results in the two studies were surprisingly different. For example, in their study 24 WPI corresponded to 75% of complete disease progression, with less than 300 DEGs, whereas the 18 WPI timepoint of our study corresponded to 80% of disease progression, with thousands of DEGs, many with |L2FC| >> 1. Furthermore, we detected over 200 DEGs at 10 WPI representing 44% disease progression, while they detected no DEGs at 16 WPI, representing 50% disease progression. Last, they detected no changes in astrocytes and microglia until late in disease, and none in neurons until mice were near death. One explanation for the discrepancy is the application of a |L2FC| > 1 cutoff in their study, which we did not apply since we expected only small changes in mild disease stages. We consider it likely that mRNAs changing less than 2-fold can have substantial biological impact. For example, given how abundant they are in the first place, it is easy to envision how the coordinated reduction of ribosomes and mitochondria in astrocytes at 10 WPI may impact on cellular metabolism, or as discussed earlier, intracellular crowding. Additional explanations for the different outcomes between the studies are technical in nature, including different mouse genetic backgrounds, different treatments of RNA (e.g., the application of RNase in their procedures), different transgenic mouse lines, etc. All explanations ought to be considered when comparing these studies.

### Outlook

With these results in hand, it will now be interesting to study additional, early time points, especially considering the drastic changes in gene expression at 18 WPI, as well as selected brain regions to further study localized subgroups of astrocytes, GABAergic and glutamatergic neurons. Furthermore, in addition to actively translated mRNAs, other levels of gene expression including miRNAs and lncRNAs as well as epigenetic features could be analyzed from specific cell types using the Tagger mouse line, analogous to the RiboTag system, but expressing four tagging modalities (Kaczmarczyk et al., 2019). Studying multiple levels of gene expression in a cell-type-specific manner will greatly enhance our understanding of disease mechanisms and may point towards therapeutic targets. The work described here, validating such a system in our mouse model in the context of an EEG-based early biomarker of disease progression, is an important first step in this direction.

## Materials and Methods

### Mouse lines and husbandry

Mice were housed in individual ventilated cages (22°C; 60% humidity) on a standard 12h light / 12h dark cycle and had access to food and water ad libitum. RiboTag mice (Jax #11029; (Sanz et al., 2009)) were crossed to cell-type-specific Cre driver lines: Vglut2-Cre (Jax #016963; (Vong et al., 2011), Gad2-Cre (Jax #010802; (Taniguchi et al., 2011), SST-Cre; Jax #013044, PV-Cre (Jax #008069; (Hippenmeyer et al., 2005)) and Cx43-CreER^T^ a gift from Martin Theis, (Kretz et al., 2003). To induce Cre activity in Rtag/Cnx43-CreER^T^ mice, starting one week before sacrifice, Tamoxifen (T5648, Sigma-Aldrich) prepared as 10mg/ml in 10% ethanol / 90% sterile sunflower oil, was intraperitoneally injected, 10µl/g body weight, once per day for three days. Experimental mice were heterozygous for both RiboTag and Cre recombinase. For each condition (5 RiboTag/Cre combinations, injected with 20µl 0.1% RML or NBH, sacrificed at 10 or 18 WPI) four mice were used (total n=80). Mice were genotyped for the presence of Cre and the RiboTag allele using an established PCR protocol (Sanz et al., 2009). Prior to intercrossing, each Cre and RiboTag mouse line was bred to S4 for at least 8 generations and were verified to be at least 99% 129S4 (Kaczmarczyk et al., 2021) by a commercial high throughput single nucleotide polymorphism service (Envigo/Harlan).

### NBH/RML prion injections

The RML prion homogenate used in these experiments was originally acquired from Gregory Raymond (Rocky Mountain Labs). It was injected into 129S4 mice at the location where this line originated, the Whitehead Institute for Biomedical Research, Cambridge, Massachusetts, resulting in an incubation period to terminal disease of 22.8 weeks. Brain homogenates prepared from these RML-infected 129S4 mice resulted in an incubation period to terminal disease of 22.6 weeks and were the source of RML for the EEG, JL and RiboTag experiments reported here. All three experiments included a common injection procedure: 20µl 0.1% brain homogenate from normal brain (NBH) or RML infected mice was injected into the right brain hemisphere at the bregmatic suture using hand-held syringes with needle guides to control injection depth of approximately 3.5 mm.

### EEG and sleep recordings

For recordings of EEG, EMG, body temperature, and locomotor activity, ten male wild type 129S4/J (Jax #009104) mice were bred in-house and injected with NBH (n=5) or RML (n=5). Only males were used to reduce the variability of sleep parameters influenced by sex (Mong et al., 2011; Mong & Cusmano, 2016). They were injected at 1 year of age since the study was done in parallel with a similar study of inherited prion diseases that required older mice (manuscript in preparation). Mice injected at this age die from RML only 5% faster than young mice (Sorce et al., 2020). Surgical procedure: After at least twelve and a maximum of 22 d incubation time, mice were implanted intraperitoneally with F20-EET transmitters (Channel bandwidth 1-50Hz, Data Sciences International (DSI)) under isoflurane anesthesia. EEG leads were routed subcutaneously to the skull, placed epidurally above the left frontal cortex (AP: 1.5, ML: 1.5 mm from bregma, negative lead) and the right parietal cortex (AP: -2.5, ML: 2.0, positive lead) and fixed in place with dental acrylic. EMG leads were anchored in the neck muscles. Mice were allowed at least two weeks of recovery before recordings. Data acquisition and analysis: Undisturbed 24 h baseline recordings were performed in the home cages. Mice were kept in individual cages but could hear, smell and see at least one other mouse of the same experiment. The cages were placed in ventilated cabinets with a 12h light / 12h dark cycle. Sleep scoring and analysis was done as reported before (Dittrich, Morairty, Warrier, & Kilduff, 2015; Morairty et al., 2013; Parks et al., 2016). EEG and EMG were recorded via telemetry using DQ ART software (DSI). Sampling frequencies were 500Hz. EEG low-pass filter cut off was 100Hz (in addition to the 1Hz high pass and 50Hz low pass anti-aliasing filtering built into the transmitter). EEG and EMG recordings were scored in 10s epochs as wake, rapid eye movement sleep or non-rapid eye movement sleep by an expert scorer who examined the recordings visually using NeuroScore 3.0 software (DSI). EEG spectra were analyzed with a fast Fourier transform algorithm using a Hanning Window without overlap (NeuroScore) on all epochs without stage transition or artifact. For direct comparisons of EEG power spectra, power was expressed as relative power, i.e., each frequency bin (0.122Hz) was divided by the sum of the values between 0 and 50 Hz. Relative theta power was calculated as the power between 5 and 10 Hz (summed values of the respective frequency bins) divided by the sum of the values between 0 and 50 Hz.

### Jet lag

The jet lag (JL) experiment was designed to study 20 male mice in parallel, our maximum recording capacity, as follows: n = 7 each for RML + JL and RML undisturbed control; n=3 each for NBH + JL and NBH + undisturbed control. To ensure the loss of a mouse from the injection or implant procedures would not compromise the experiment, NBH and RML were each injected into an extra mouse. While they were not needed for the EEG study, they were both subjected to JL and included in the histology study. Since it was not feasible to obtain such a large cohort of males by breeding in our facility, we imported C57Bl/6JRj male mice at approximately 3 months of age from Janvier in France. Following injections with NBH or RML, mice were kept in two identical microisolator cabinets, one for the jet lag and the other for the undisturbed control group, with 12-hour light cycles initially set for lights on at 9:00. Two weeks after injections surgeries were performed to implant the telemetric recording devices, as done above. The first 24h baseline recordings were done at 5 and 6 WPI. Immediately after the 6 WPI recordings, lights for the JL group were subjected to a phase advance (PA) whereby the lights turn on and off 6 hours earlier in the day. After one week of habituation to the advanced Zeitgeber, EEG was recorded again, and the cycle of PA followed by EEG 1 week later was repeated for a total of 6 PAs. PAs were always accomplished by shortening the light phase. Following the final PA at 11 WPI, additional EEG recordings were made at 12, 13 and 14 WPI. All 24 h recordings were started at the same clock time irrespective of light status, keeping the distance between recordings constant. Since the most rigorous analysis of EEG data requires scoring by a human expert, and the scoring involves inspection of data at a resolution of 10 second bins, it was unfeasible to score all data. We expected that at 12 WPI there might be residual effects from the previous PA affecting EEG, independent of disease. However, we thought that by 13 WPI, the immediate effects of PA would have passed, and we would be able to measure the lasting impact of JL on theta, and thus focused on these data for the analysis. Mice were sacrificed immediately after the last recording and brains were processed as described above. JL and undisturbed control brains were embedded in blocks together to ensure equal processing and staining and to facilitate microscopic evaluation.

### Tissue preparation

For gene expression studies 2-4-month-old RiboTag mice were injected, 4 mice each for NBH and RML. Only females were used for Gad2, PV, and Cx43 lines; both sexes were used in nearly equal proportion for vGlut2 and SST lines. RiboTag mice were sacrificed at 10 or 18 WPI by CO2 asphyxiation. Brains were removed from the skull, the olfactory bulbs were removed and discarded, the brain stem was cut at the level of the most caudal tip of the cerebellum, the two hemispheres were separated; one was snap frozen for mRNA isolation and the other formalin fixed for IHC. Prion infectivity in formalin fixed hemispheres was inactivated with formic acid prior to embedding in paraffin cassettes. Histology samples from mice injected with NBH were treated the same way in parallel.

### RiboTag translational profiling

Each frozen brain hemisphere was weighed and put into a tissue grinder with a corresponding volume of polysome buffer (50mM Tri pH = 7.5, 100mM KCl, 12mM MgCl2, 1% Nonidet P-40, 1mM DTT, 1x Protease inhibitor (SigmaFast Protease Inhibitor Tablets (S8820), Sigma-Aldrich), 100U/ml RNAse inhibitor (RNAase inhibitor (N8080119), Thermo Fisher Scientific), 100µg/mL cycloheximide (Cycloheximide (C7698), Sigma-Aldrich) to prepare a 10% brain homogenate. Homogenate was prepared with help of a motorized tissue grinder (Heidolph, 600rpm, ∼30s) on ice and centrifuged (10000g, 10min, 4°C). Supernatant was transferred to a new reaction tube and used as IP input and for isolation of total RNA. For IP with magnetic beads (Dynabeads Protein G 10004D, Thermo Fisher Scientific) pre-cleared supernatant (25µl beads and 200µl supernatant, 30min, 4°C) was first incubated with anti-HA antibody 12CA5 (Roche Life Science; 200µl pre-cleared supernatant, 10µl antibody, 45min, 4°C) and this mixture then added to the magnetic beads (50µl beads, 1-2h, 4°C). Magnetic beads were washed three times with PBS (∼500µl) before use and incubation steps of IP were done on a rotator. IP samples were put on a magnetic rack and the magnetic bead pellets were washed three times with high salt buffer (50mM Tris pH = 7.5, 300mM KCl, 12mM MgCl2, 1% Nonidet P-40, 1mM DTT, 100µg/ml cycloheximide; ∼500µl). Cell-type-specific mRNA was eluted from the magnetic beads by RLT buffer supplemented with 2-mercaptoethanol from the RNeasy Mini Kit (Quiagen; 200µl; Thermomixer: 700rpm, 5-10min, RT) and afterwards isolated with this kit. For each supernatant two technical replicates of IP were done, and total RNA from the input supernatant (200µl) was isolated in parallel. Quality and quantity of immunoprecipitated mRNA and total RNA were verified by Qubit Fluorometer (Thermo Fisher Scientific) and Agilent 2100 Bioanalyzer (Agilent Technologies).

### RNA-seq library preparation and RNA-sequencing

300ng of total RNA or 150ng of immunoprecipitated mRNA was used for RNA-sequencing. Each sample was checked for amount and quality using a Nanodrop 2000 (Thermo Fisher Scientific) and Agilent 2100 Bioanalyzer. For each condition (5 Cre/Rtag lines, injected with RML or NBH, sacrificed at 10 or 18 WPI) we used four individual samples (exceptions: SST RML 10 WPI: n = 3; PV NBH 10 WPI: n = 3; Astro RML 18 WPI: n = 2). RNA was converted to cDNA using the Transcriptor High Fidelity cDNA Synthesis Kit (Roche Aplied Science). RNA-sequencing libraries were prepared using the TruSeq RNA Sample Preparation 2 Kit (Illumina). The library quality was checked using an Agilent 2100 Bioanalyzer and concentration was measured by a Qubit dsDNA HS Assay Kit (Thermo Fisher Scientific) and adjusted to 2nM before sequencing (single end, 50 bp) on a HiSeq 2000 Sequencer (Illumina) using TruSeq SR Cluster Kit 3-cBot-HS (Illumina) and TruSeq SBS Kit 3-HS (Illumina) according to the manufacturer’s instructions.

### RNA-seq data analysis

Quality assessment was based on the raw reads using the FastQC (0.10.1, Babraham Bioinformatics) quality control tool. The sequence reads (single-end, 50 bp) were aligned to the mouse reference genome (mm10) with Bowtie2 (2.0.2) (Langmead & Salzberg, 2012) using RSEM (1.2.29) (Li & Dewey, 2011) with default parameters. First, the mouse reference genome was indexed using the Ensembl annotations (84.38) with rsem-prepare-reference from RSEM software. Next, rsem-calculate-expression was used to align the reads and quantify the gene abundance. Differential expression analysis was carried out using total gene read counts with DESeq2 package (1.12.4) (Love, Huber, & Anders, 2014). Genes with less than five reads (baseMean) were filtered out and false discovery rate (padj/FDR) was recalculated with Benjamini-Hochberg procedure for the remaining genes.

### Overrepresentation analysis (ORA) and gene set enrichment analysis (GSEA)

ORA was performed with the `enricher()` function from clusterProfiler R package (Yu, Wang, Han, & He, 2012). GSEA was done using WebGestallt (Liao, Wang, Jaehnig, Shi, & Zhang, 2019). Gene sets were sourced from Molecular Signature Database 3.0 (Liberzon et al., 2011) and sets containing less than 10 or more than 500 genes were excluded from the analysis. For ORA, all annotated genes expressed in the analyzed disease time point and cell-type were used as background gene set. Redundant categories were filtered out using simplifyEnrichment R package (v.1.3.1) (<u>10.18129/B9.bioc.simplifyEnrichment</u>) and visualized using ggplot2 R package (v.3.3.5) (https://ggplot2.tidyverse.org).

### Gene network construction for disease module analysis

Protein-protein interaction network for Mus musculus was sourced from the STRING database (v. 11.0) (Szklarczyk et al., 2021). ENSEMBL protein IDS representing node names were replaced with corresponding ENTREZ gene IDs using the `getBM()` function (biomaRt package; version 2.64.3). Data were filtered for interactions with combined confidence score ≥ 0.7 and a separate graph object was created for all cell types (igraph package; version 1.2.6), including only the vertices with corresponding Wald test p-values (genes that were not expressed in either RML or NBH mice were removed). Topological characteristics of all generated networks are presented in Table S1. Interactome information was sourced from the STRING database (v. 11.0) (Szklarczyk et al., 2021). The interactome igraph object used in this study is available from GitHub (https://tinyurl.com/tcyryath).

### Seed clusters validation for disease module analysis

We have considered the genes with FDR < 0.1 as members of the seed cluster and used a network-based approach to infer cell-type-specific disease modules. For a given set of seed genes LCC size was measured with components() function (igraph R package) and compared to a reference distribution obtained by measuring the LCC size of 10,000 random node subsets of size equal to that of the original seed gene set. Randomization was done so that node selection was limited to genes with degrees comparable to those of the seed cluster. To avoid repeated selection of high degree nodes (in biological networks such nodes are rare, thus many have unique degree values), nodes within a specified degree interval were organized into bins such that there were at least 100 nodes in each bin. Random sampling of nodes was then performed from bins containing seed genes (degree-preserving randomization). Statistical significance of the observed LCC size was assessed by calculating the empirical p-value (the fraction of random sets of nodes with LCC size equal to or greater than the size of LCC formed by seed genes), and Z-scores (Z-score =(LCC_O_ - mean(LCC_R_))/sd(LCC_R_), where LCC_O_ is observed LCC LCC_R_ is random LCC and sd is standard deviation). All scripts are available from GitHub (https://tinyurl.com/5sbexp89).

### Construction and validation of cell type-specific disease modules

Candidate disease module genes were identified for selected cell types using the modified DIAMOnD algorithm, consisting of the following steps: 1) for all genes connected to any of the seed nodes, Wald test p-values (the minimum value was selected in cases where multiple Wald test p–values corresponded to the same ENTREZ gene ID) and connectivity significance were determined, 2) candidate proteins were ranked based on their respective Wald test p-values and connectivity significance (ties within individual rankings were resolved using fractional ranking), 3) Individual rankings were combined into a single score given as the sum of the inverted ranks. 4) the gene with the highest combined score was included into the disease module and became a member of the new seed cluster, 6) steps 1-5 were repeated 500 times. The threshold for the total number of module genes was set as the iteration at which the number of genes with Wald test p-value < 0.01 in the module reached a plateau (Fig S5B). The relevance of individual thresholds was verified by determining the number of module genes sharing functional annotations with seed nodes. Annotated gene sets from Gene Ontology (GO), KEGG, Reactome and Biocarta were retrieved using the msigdbr R package (v7.4.1) and all GO terms/pathways significantly enriched within a given set of seed genes were identified using Fischer’s exact test (Benjamini–Hochberg adjusted p-value < 0.05) (Gene sets containing less than 10 and more than 500 genes were excluded from the analysis). Module genes annotated to any of the identified GO terms/pathways were considered true positives. The statistical significance of the number of true positive genes at a given iteration step was determined using Fischer’s exact test). Gene modules were validated using annotated gene sets from Gene Ontology (GO), KEGG, Reactome and Biocarta were retrieved using msigdbr R package (v. 7.4.1) and used in ORA analysis using clusterProfiler R package. Genes annotated to any of the identified GO terms/pathways were included in the validation gene set. The statistical significance of the number of functionally related module genes at a given iteration step was determined using one-sided Fisher’s exact test.

### Data availability

The raw and preprocessed data discussed in this publication have been deposited in NCBI’s Gene Expression Omnibus (Edgar, Domrachev, & Lash, 2002) and are accessible through GEO Series accession number GSE189527 (https://www.ncbi.nlm.nih.gov/geo/query/acc.cgi?acc=GSE189527).

## Acknowledgments

This work was supported by the Knut and Alice Wallenberg foundation, Deutsche Forschungsgemeinschaft (DI 1718/3-1 to LD), Helmholtz-Alberta Initiative- Neurodegenerative Disease Research (HAI-NDR SO-083 to WSJ), and internal funding from the DZNE.

---

**Figure S1.**
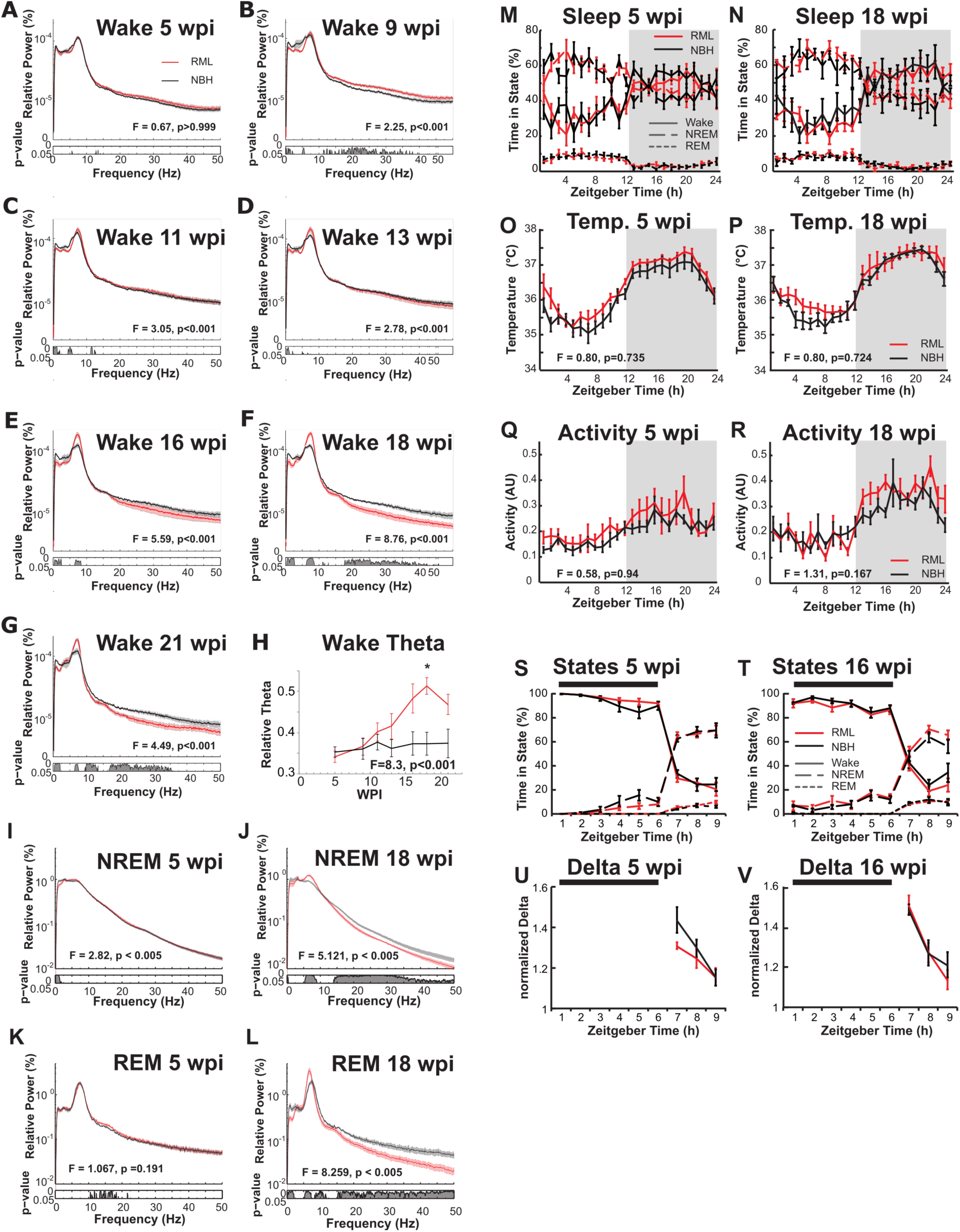
Characterization of EEG and sleep features in RML. (A-L) Frequency power spectra from serial EEG. (**A-G**) Examples of frequency power spectra during wake, acquired serially as disease progressed. The power in each frequency bin is expressed as percentage of the cumulative power of all frequencies (0-50 Hz). Curves depict group averages, shaded areas depict SEM. The degrees of freedom are 409 and 3272. The p-values for post hoc uncorrected bin-by-bin t-tests are indicated below the spectra. The average wake power spectra up to 11 WPI (**C**) shows no difference between the groups but at 13 WPI (**D**) differences begin to emerge and at 18 WPI (**F**) RML mice show a significant increase in relative power in the theta range (5-10 Hz) and a decrease in relative power in lower and higher frequencies. A summary of data in panels A-G is depicted in H, also shown in Fig 1C. A similar increase in theta at 18 WPI was also seen for NREM (**J**) and REM (**L**), but not at 5 WPI (**I**, **K**). <u>(**M-R**) **Baseline** vigilance states, temperature and activity in RML and control mice, relative to</u> **<u>time</u>.** (**M**, **N**) Average Wake, NREM and REM time during 24 h undisturbed recording. The shaded area indicates the 12 h lights-off period. Mixed model ANOVA did not reveal significant main effects for group (RML or NBH) nor interactions between group and Zeitgeber Time (ZT, hourly bins) for any vigilance state at either 5 WPI (M) or 18 WPI (N). (**O**, **P**) Average intraperitoneal temperature across 24 h undisturbed recording. The average temperature did not differ between groups (no main effect found by ANOVA), and the course of temperature over time was also not affected by treatment (no interaction of temperature and ZT) at 5 and 18 WPI (O and P, respectively). The ANOVA result for the interaction is given at the bottom of the panel. (**Q**, **R**) Average locomotor activity across 24 h undisturbed recording. Activity levels did not differ between groups (no main effect), and there was no interaction between group and ZT (indicated at bottom of graphs) for 5 or 18 WPI (Q and R, respectively). Degrees of freedom for ANOVAs were 1, 8 for main effects of treatment group and 23, 184 for interaction effects of treatment group and ZT (hourly bins). <u>(**S-V**) **Response to**</u> **<u>6 h sleep deprivation in RML and control mice</u> (S**, **T**) Vigilance states (wake, NREM, REM) across 6 h of total sleep deprivation by the gentle handling method and the subsequent 3 h of recovery sleep opportunity at 5 and 16 WPI (18 WPI was not tested). The sleep deprivation period is indicated by the horizontal black bar at the top of each panel. T-tests of the first hour of recovery (ZT7) did not show significant differences for either state at 5 WPI (S, p > 0.21) or at 16 WPI (T, p > 0.48). (**U**, **V**) Average NREM EEG delta power (0.5-4Hz) during sleep deprivation and recovery, normalized by the average NREM delta power of the corresponding undisturbed baseline recording. Both groups show the expected increased delta power (values higher than 1) during recovery from sleep deprivation. However, a T-test of the first hour of recovery (ZT7) did not show a significant difference between groups at either 5 WPI (U, p = 0.078), or 16 WPI (V, p = 0.669).

**Figure S2.**
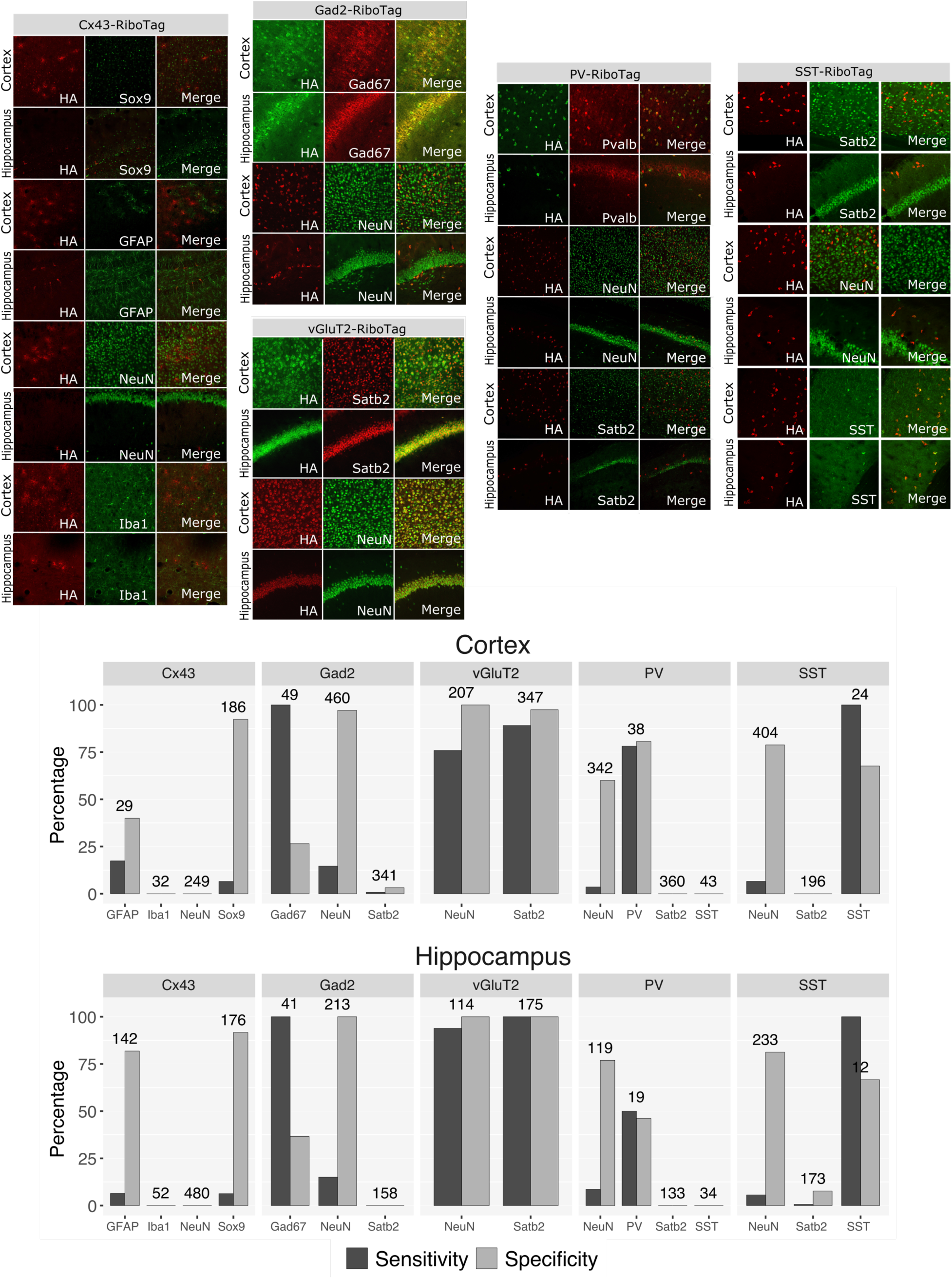
Immunofluorescence showing RiboTag expression specificity and sensitivity. (**A**) Confocal images of double immunofluorescence labeling of HA (RiboTag) and representative histological cell-type markers. (**B**) Bar plots showing the numbers of cells immunopositive for HA and appropriate histochemical markers. Sensitivity is the percentage of cell profiles positive for the marker that is also positive for HA. Specificity is the percentage of cell profiles positive for HA that is also positive for the marker.

**Figure S3.**
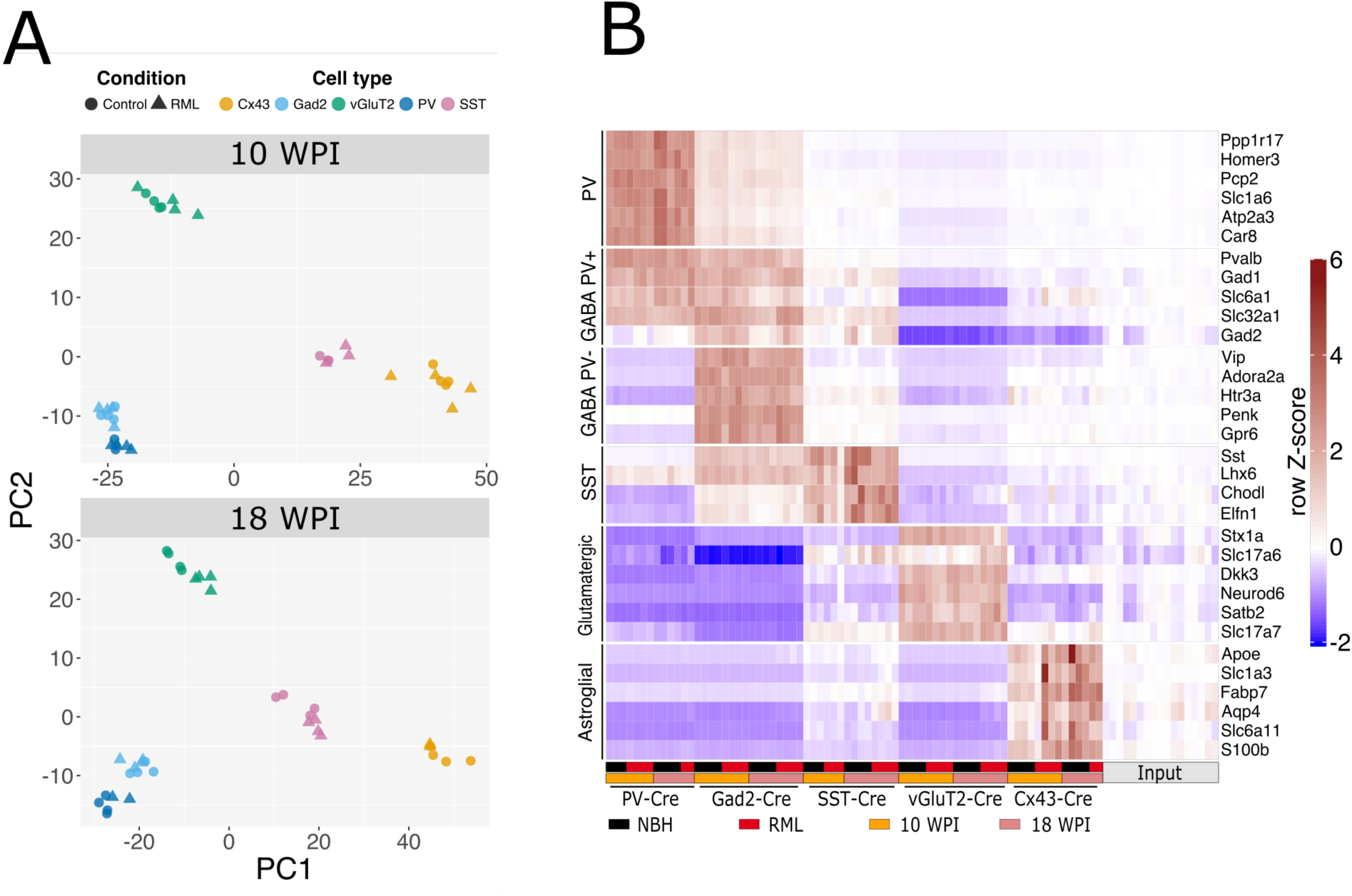
Translatome sequencing shows RiboTag expression specificity and sensitivity. (**A**) Principal Component Analysis (PCA) of RiboTag data from all cell types and time points analyzed. PCA was done on variance-stabilized normalized counts (`vst()` transformation using DEseq2 R package). Clustering was dominated by differences between cell types rather than disease states. (**B**) Heatmap showing RiboTag specificity. Each row corresponds to a cell type marker for one of the cell types listed on the left side of the heatmap. Each column represents one biological replicate (one mouse), and the specification of the sample is encoded by the colored legend below the heatmap. Input samples are total RNA collected specifically for this analysis for comparative purposes. Z-score for each row was calculated so that the mean input level for each row was set to 0 (`Z=(x–row_mean`_input_`)/row_SD`, where `row_SD` is row standard deviation and `x` is the TPM value).

**Figure S4.**
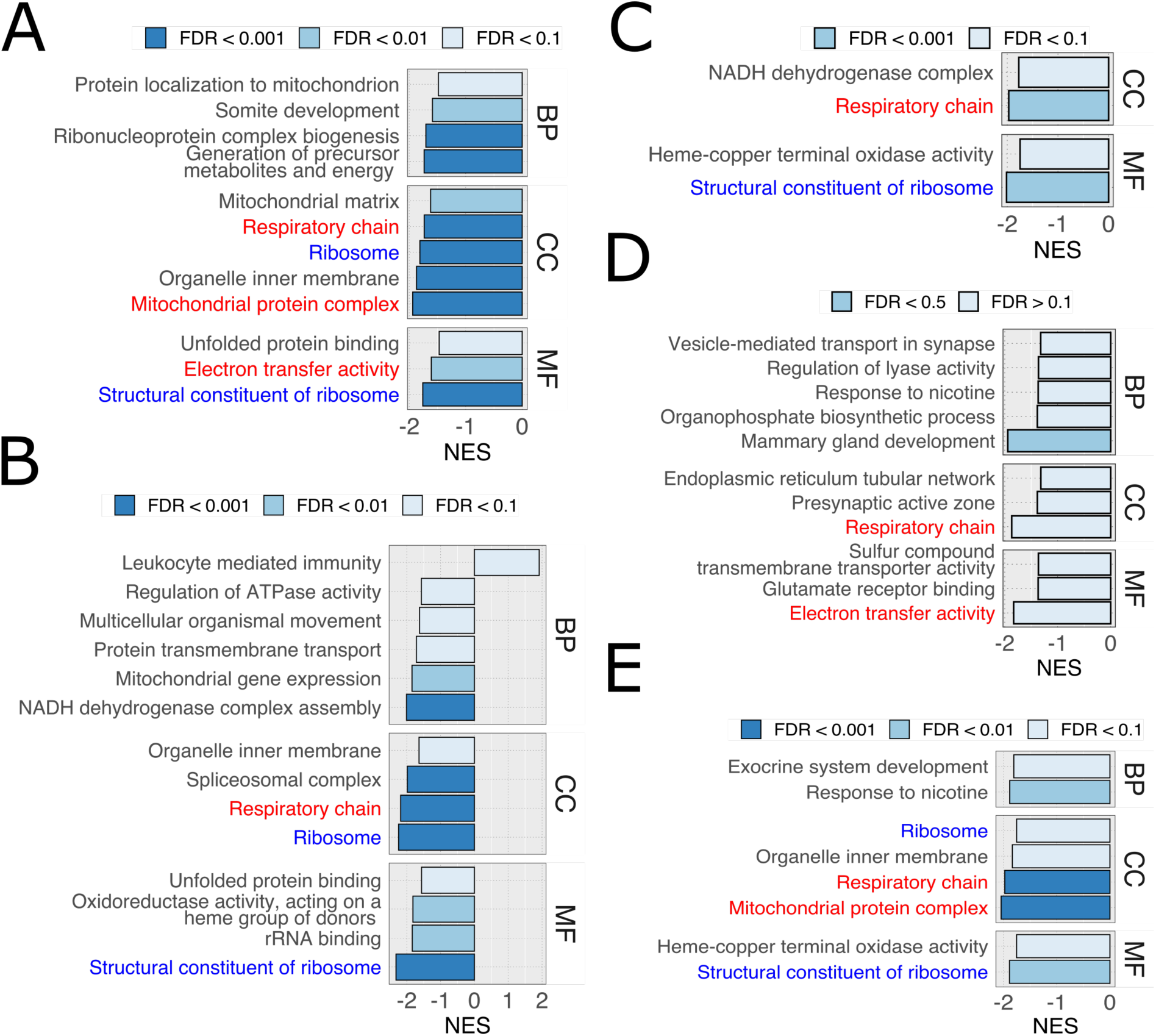
Gene Set Enrichment Analysis (GSEA) of 18 WPI samples. (**A**-**E**) GSEA of DEGs in (respectively) astrocytes, glutamatergic neurons, SST neurons, PV neurons and GABAergic neurons at 18 WPI. In case of PV neurons (D), no terms enriched at FDR < 0.1 were found and therefore terms > 0.1 were shown. Categories with GSEA FDR < 0.1 were shown after removing similar terms based on semantic similarity using GOSemSim R package (see methods for details). Categories related to terminal oxidation (highlighted in red) and translation (highlighted in blue) were amongst the most enriched, and were common across multiple analyzed cell types, suggesting that the 18 WPI timepoint is marked by canonical changes to mitochondria and ribosomes, typical for neurodegenerative diseases.

**Figure S5.**
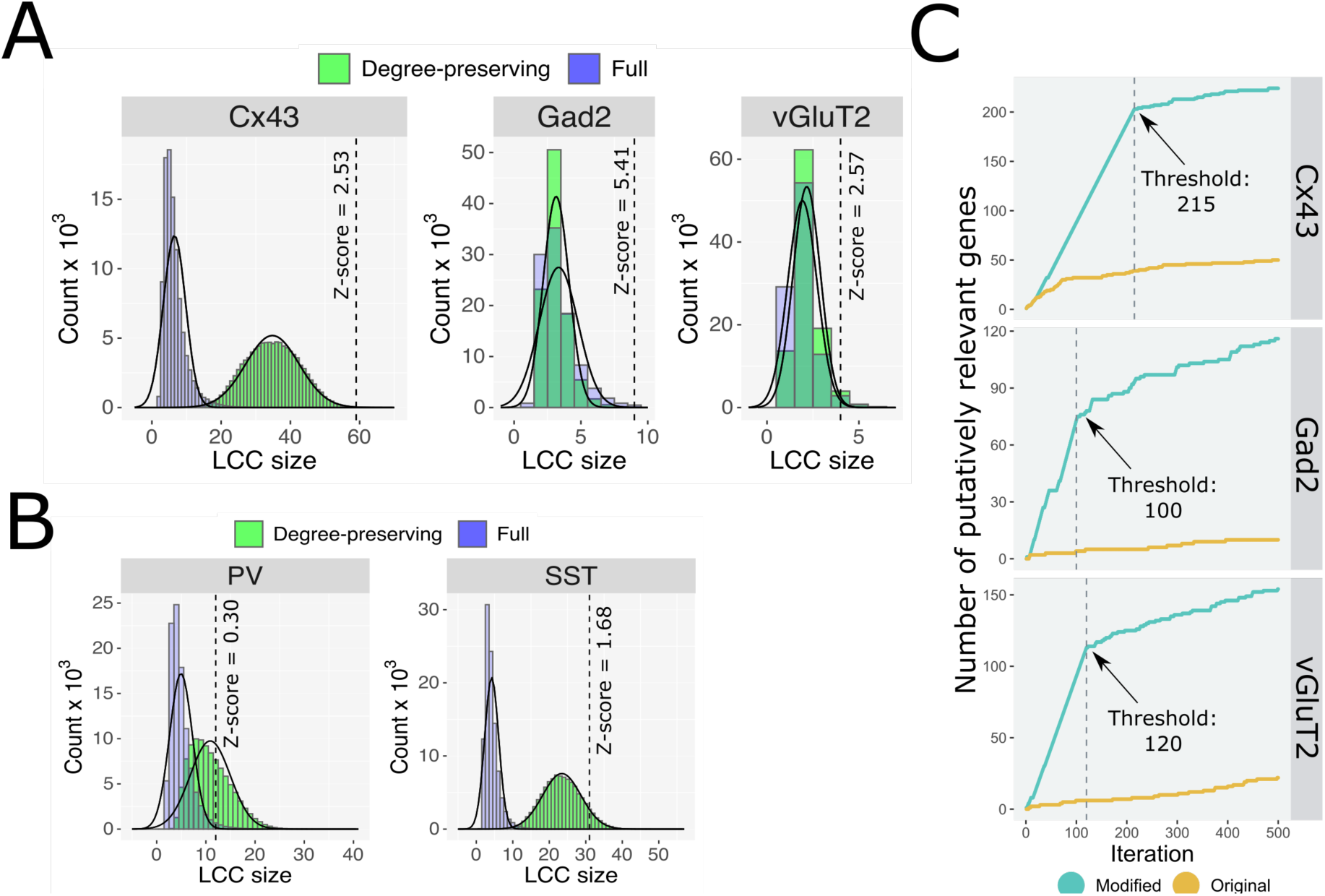
LCC size compared to randomly generated control LCCs. (**A**, **B**) LCCs were calculated for each cell type and compared to randomly generated LCCs in which random genes were limited to those with similar connectivity as the test genes (degree-preserving) or were full random. (**A**) For affected cells (Cx43, Gad2 and vGluT2), LCC sizes were larger (indicated by dotted lines) than control LCCs computed using either method. (**B**) PV and SST cells had LCCs that were not larger than the degree-preserving controls. (**C)** Determination of lists sizes was based on the breakpoint threshold resulting from repeated additions of one gene at each iteration.

**Figure S6.**
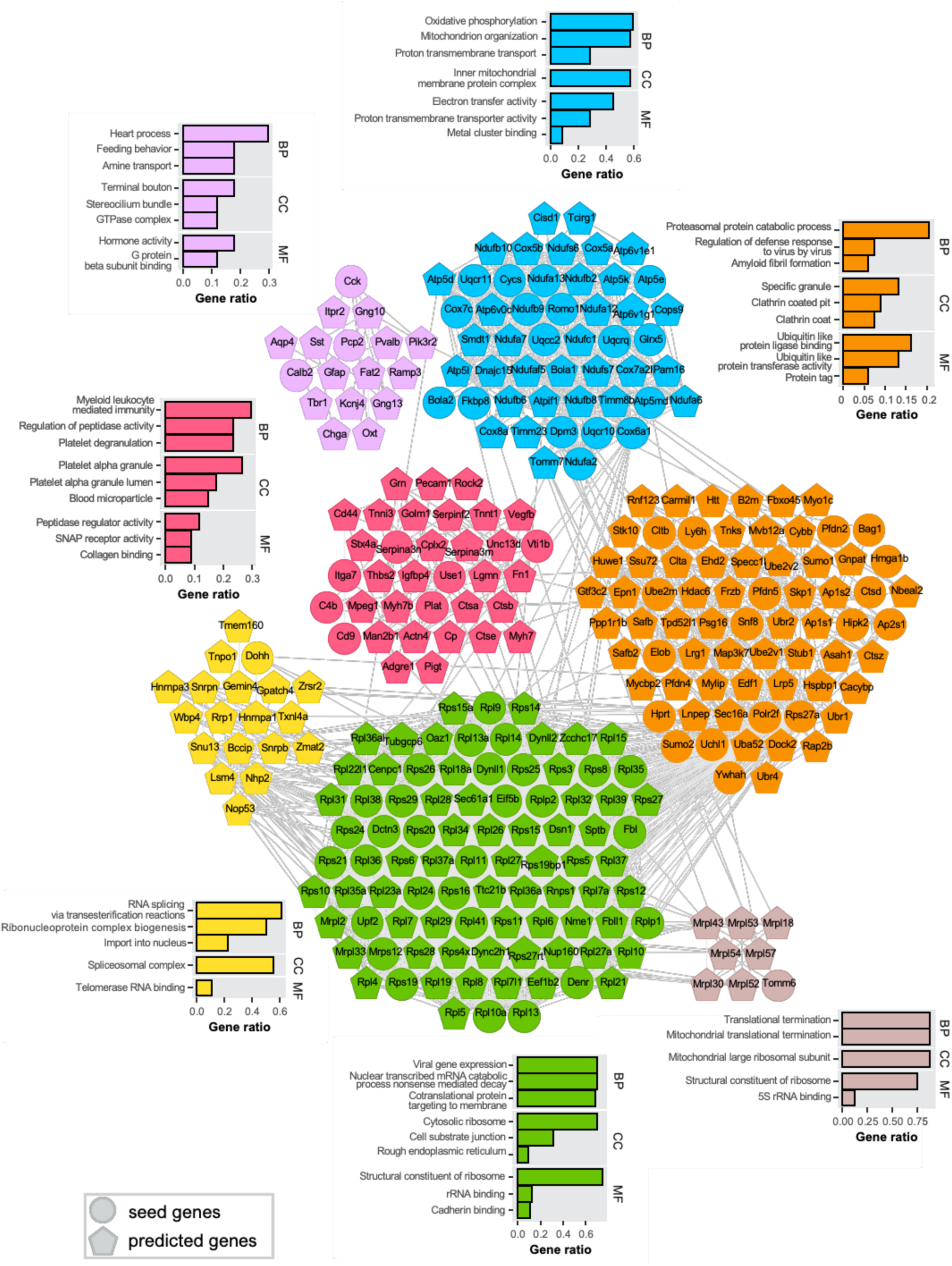
Overrepresentation analysis of astrocytic (RiboTag::Cx43-Cre) DIAMOnD module. Colors represent fast-greedy clustering of genes within the module.

**Figure S7.**
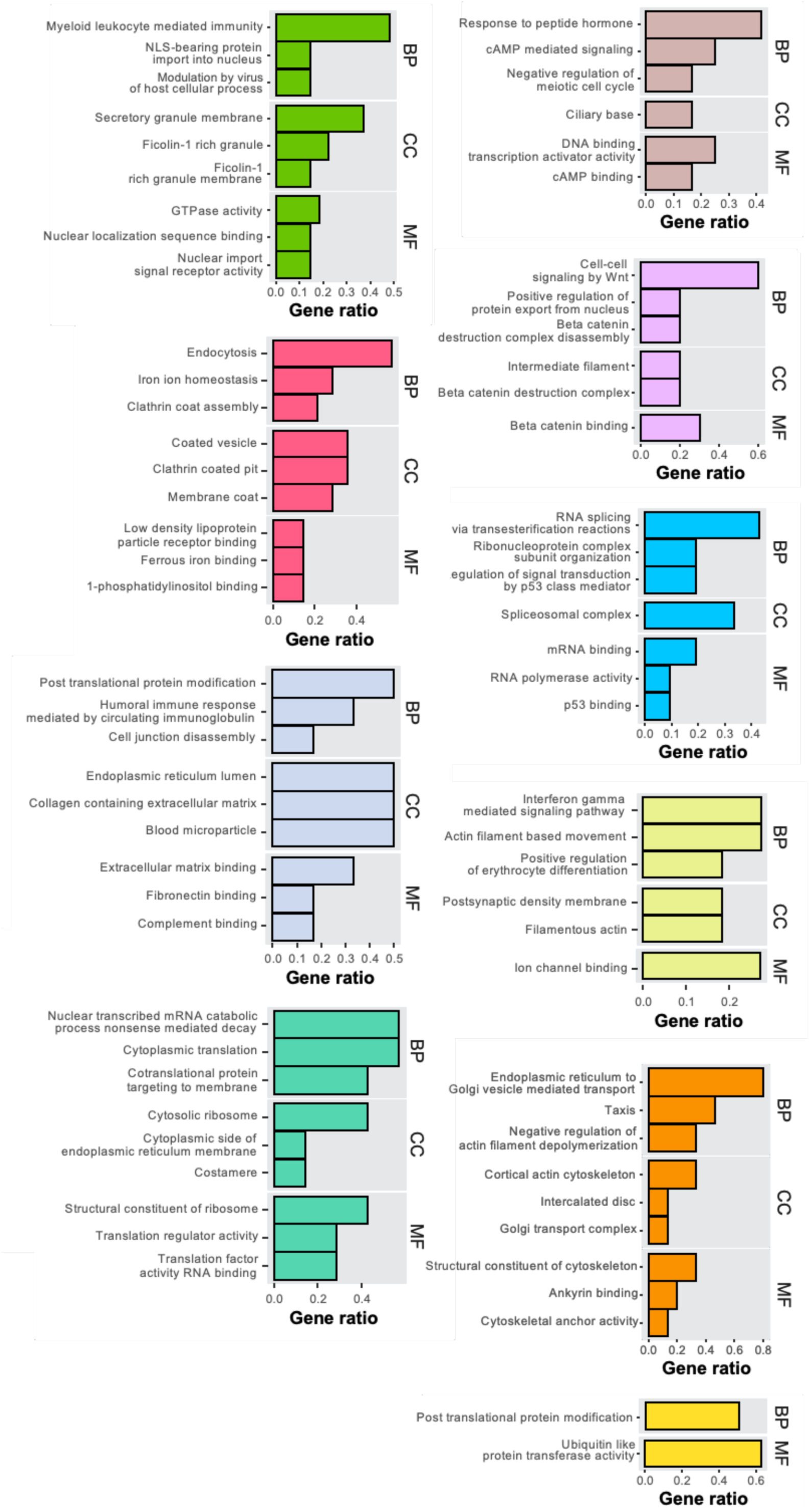
Overrepresentation analysis of glutamatergic (RiboTag::vGluT2-Cre) DIAMOnD module. Colors represent fast-greedy clustering of genes within the module.

**Figure S8.**
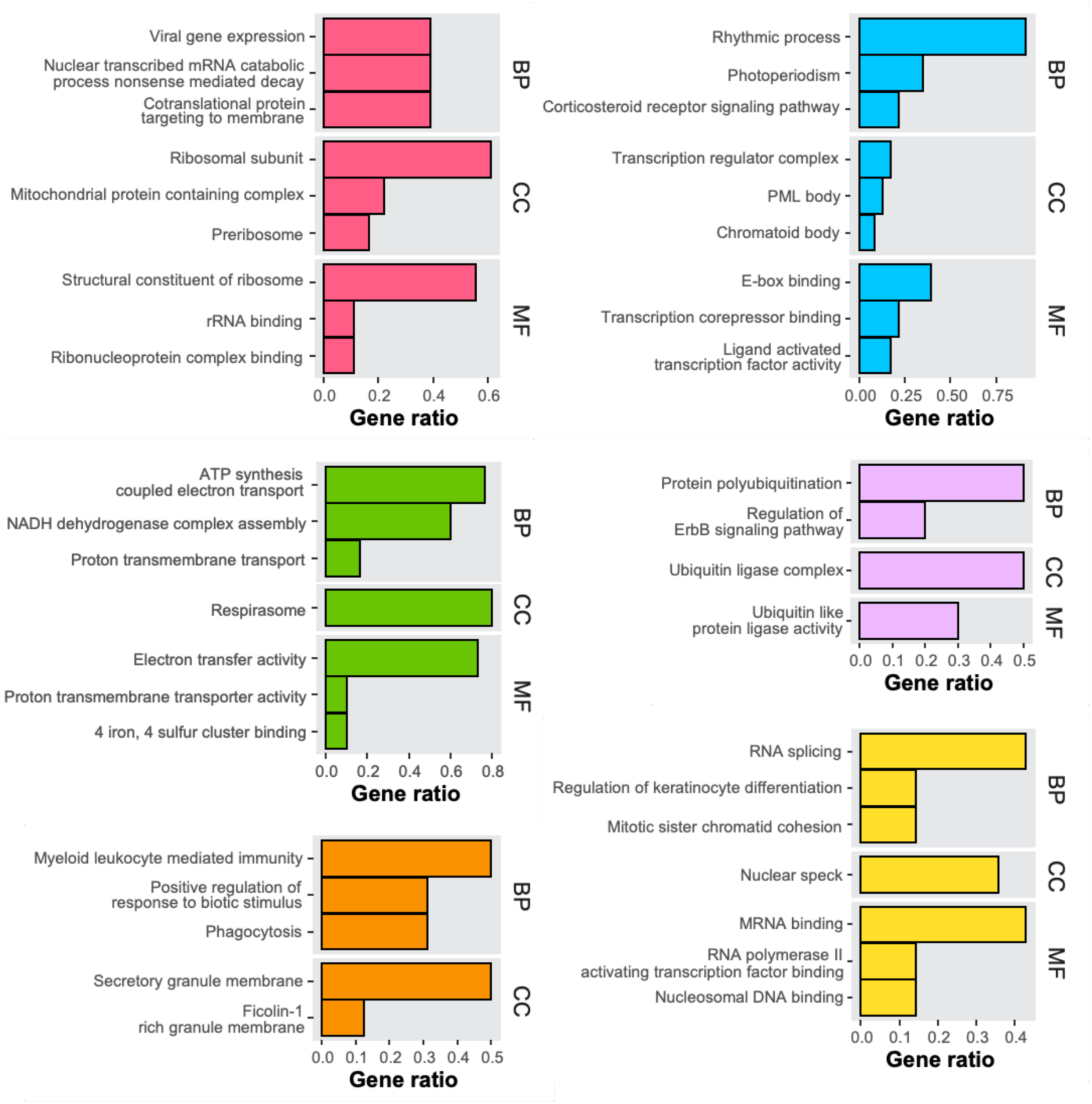
Overrepresentation analysis of GABAergic (RiboTag::Gad2-Cre) DIAMOnD module. Colors represent fast-greedy clustering of genes within the module.

**Figure S9.**
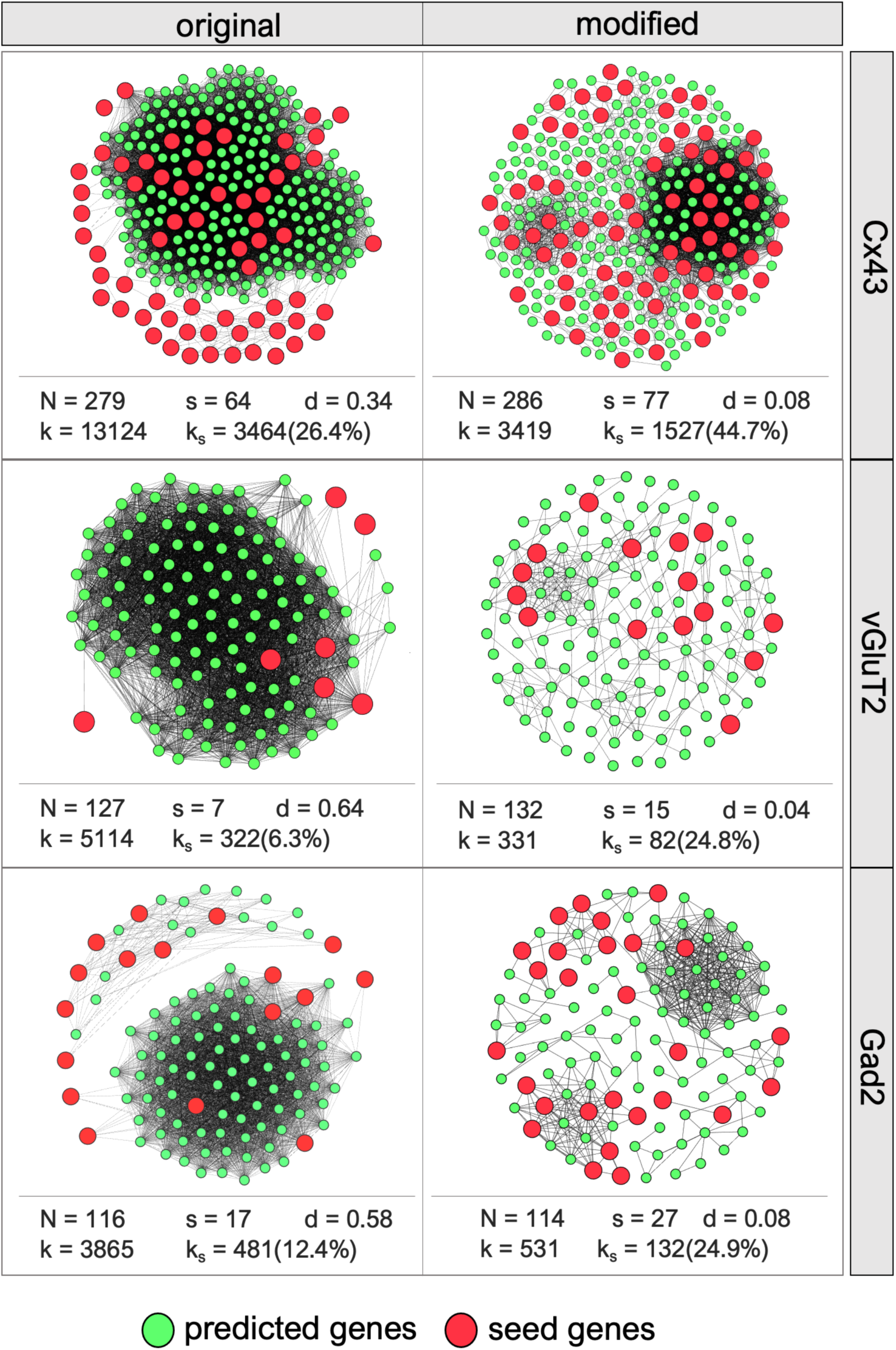
Topological characteristics of disease modules obtained with original (left) and modified (right) DIAMOnD algorithm. N - number of nodes, s - number of seed nodes in the final module, d - edge density, k - number of edges, ks - number of edges including predicted and seed nodes (ks). The modules produced by the modified algorithm had more seed genes and a higher percentage of connections between initial seed genes and predicted genes than those obtained from the original algorithm.

**Table S1.**
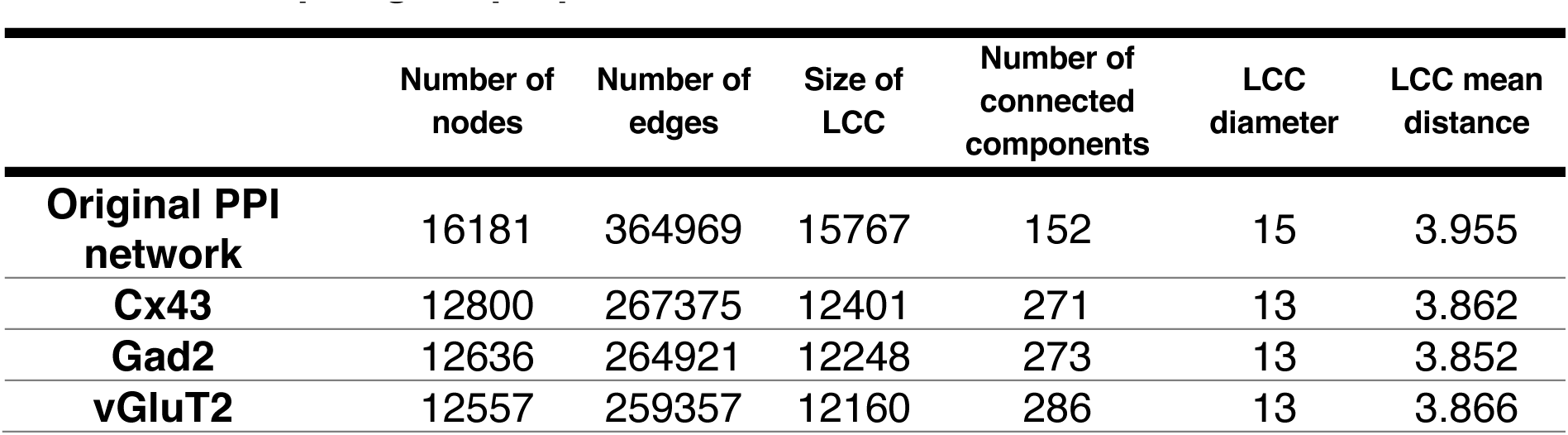
Topological characteristics of all generated networks are presented. In contrast to a previous report (Magger, Waldman, Ruppin, & Sharan, 2012), we observed that node removal does not have a major impact on network connectivity. As indicated by the number of connected components, filtered networks are more fragmented in comparison to the generic network. However, most of the genes constitute a single connected component (96.9% on average). The diameter and mean shortest distance did not increase with node removal, suggesting that the “small world” property of the original network was preserved.

|  | Number of nodes | Number of edges | Size of LCC | Number of connected components | LCC diameter | LCC mean distance |
| --- | --- | --- | --- | --- | --- | --- |
| **Original PPI network** | 16181 | 364969 | 15767 | 152 | 15 | 3.955 |
| **Cx43** | 12800 | 267375 | 12401 | 271 | 13 | 3.862 |
| **Gad2** | 12636 | 264921 | 12248 | 273 | 13 | 3.852 |
| **vGluT2** | 12557 | 259357 | 12160 | 286 | 13 | 3.866 |

**Table S2.** Seed gene information and validation results. Only a fraction of seed genes have corresponding vertices in the PPI network. Most of the seeds occupy neighboring interactome positions, as indicated by the significance of their LCC size (empirical p-value < 0.05). Note that sizes of LCCs formed by seeds were compared to a reference distribution which was obtained by measuring the LCC size of random genes with degrees similar to the ones of the initial seed cluster. Thus, the observed local clustering cannot be attributed solely to the network’s structural characteristics, but points to the network localization of the seed genes themselves, highlighting their relevance in marking the disease-related interactome community.

|  | Number of seed genes | Number of seed genes in the PPI network | Observed LCC size | Empirical p-value |
| --- | --- | --- | --- | --- |
| **Cx43** | 137 | 128 | 55 | 0.0048 |
| **Gad2** | 83 | 69 | 9 | 0.0014 |
| **vGluT2** | 38 | 33 | 4 | 0.0426 |

## Supplementary methods

### Characterization of module topology

Topological characteristics of generated graphs for the original and modified DIAMOnD methods were determined with the following functions of the igraph R package: `gorder()` (number of nodes), `gsize()` (number of edges), components (number and size of connected components), `diameter()` (size of the longest of all shortest paths in the module), `mean_distance()` (average length of all pairwise shortest distances), `edge_density()` (the number of edges divided by the number of possible edges). The genes constituting individual disease modules constructed with the modified DIAMOnD algorithm were assigned to unique communities using the fast greedy modularity optimization algorithm (`cluster_fast_greedy()` function in igraph package). Each module gene set with size ≥10 was then tested for enrichment with GO terms. The term with the lowest Benjamini Hochberg adjusted p-value was chosen as a representative term of each community.

### Cell counting to estimate RiboTag expression fidelity

To assess the overlap between cells expressing the HA-tag and cells immunopositive for select markers, we counted cell profiles in hippocampus and cortex. A picture of 320×320 µm was taken with a confocal microscope (LSM 700, Zeiss) using a 20x objective and all labelled cell profiles counted. For the hippocampus, we selected CA1. In cases where CA1 showed no or very few labelled cells, we analyzed the hilus region. For the cortex, we selected the deep layers right above CA1. If the number of HA+ cells was low, an additional second or third picture was taken, so the minimal number of HA+ cells in one analysis was 10. For images selected for publication, brightness and contrast were enhanced using ImageJ.

### Modification of the DIAMOnD method

First, we constructed a PPI network using the STRING database (Szklarczyk et al., 2021). We considered all interactions in the filtered networks, regardless of the species in which they were primarily described. Although we didn’t choose to further trim the interactome data, we observed that only a fraction of seed genes has corresponding vertices in the PPI network which may serve as proof of its incompleteness (Table S2). However, we found that most of the seeds occupy neighboring interactome positions (Table S2). With consideration of the shortcomings of the PPI networks, we sought to develop an algorithm relying on the connectivity patterns of seed genes as well as on the information contained in gene expression data. We chose to modify the existing DIAMOnD algorithm (Ghiassian et al., 2015). For every candidate gene, we considered the significance of the change in its expression levels (Wald test p-value from DEseq2) and connectivity significance as a measure of the gene’s topological relevance. Specifically, we have identified all seed gene neighbors and ranked them according to their respective connectivity significance p-values and differential expression (Wald test) p-values. The individual rankings were then combined into a single integrated score. The gene with the highest score was included in the growing disease module. As in the original algorithm, each identified gene was considered a member of the seed cluster in the following iteration (see also “Construction and validation of cell type-specific disease modules” in the main methods). This approach prevents omissions of potential genes of interest that could simply be excluded due to the following possibilities: 1) conservative criteria selection of DEGs based solely on an arbitrary significance threshold, 2) changes to gene expression being too subtle to be detected due to low statistical power inherent in settings with a large number of variables and a relatively low number of biological replicates, 3) subthreshold changes of expression of several genes belonging to a functional set, but having a significant cumulative effect on the network 4) changes to certain genes not being reflected by RNA levels (e.g., protein stability affected instead) or 5) changes involving a different regulatory modality (e.g., phosphorylation). All algorithms used are available on GitHub (https://tinyurl.com/5sbexp89).

### Validation of the DIAMOnD method

We integrated the interactome and RiboTag data sets using a modification of the DIAMOnD method (Ghiassian et al., 2015) and sought to identify putative disease modules specific to each cell type in early pre-symptomatic disease stage (10 WPI). To compare the performance of the default and the modified DIAMOnD version, we first specified a second, less stringent arbitrary significance threshold (Wald test p-value from DEseq2 < 0.01) to define a set of genes that are likely to be differentially expressed but do not meet the seed cluster inclusion criteria and consider them as part of a module validation cluster (putatively regulated genes). The rationale for this approach was that if DEGs are to be primarily found at the top of a gene list ranked according to differential expression p-values, a disease module should be enriched in genes from the top of the p-value ranking. Figure S5C shows that more putatively regulated genes were found by the modified diamond method in 500 iterations. Moreover, the rate at which new putatively regulated genes were added to the module was higher up to a threshold (Fig 5C) that was used as the iteration cutoff for formulating the final modules.

### Cell counting to estimate RiboTag expression fidelity

To assess the overlap between cells expressing the HA-tag and cells immunopositive for select markers, we counted cell profiles in hippocampus and cortex. A picture of 320×320 µm was taken with a confocal microscope (LSM 700, Zeiss) using a 20x objective and all labelled cell profiles counted. For the hippocampus, we selected CA1. In cases where CA1 showed no or very few labelled cells, we analyzed the hilus region. For the cortex, we selected the deep layers right above CA1. If the number of HA+ cells was low, an additional second or third picture was taken, so the minimal number of HA+ cells in one analysis was 10. For images selected for publication, brightness and contrast were enhanced using ImageJ.

### Immunohistochemistry

Formalin fixed brain tissues were treated with 98% formic acid to reduce prion infectivity, and then post-fixed for at least four more days in formalin. Brains were then embedded in paraffin with each cassette containing both NBH and RML injected brains, ensuring that controls and diseased samples were stained identically. The necessity of formic acid pretreatment renders the brain samples in a fragile state that is most successfully prepared with paraffin sectioning. These methods result in high autofluorescence and reduce the usefulness of the samples for immunofluorescent studies. We therefore employed chromogenic staining methods for all scrapie-infected and control samples. Cassettes were cut into 4µm thick sections. Sections were dewaxed in xylene and rehydrated in graded dilutions of ethanol (each 5min). Sections stained for PrP aggregates were further treated with 98% formic acid (3 min). This step was excluded for all other stainings. Epitope retrieval was performed with a steamer in 0.01 M citrate buffer (PrP: pH = 6; Iba1, GFAP: pH = 8; 30min). Endogenous peroxidase was removed with H2O2 treatment (3%, 5 min). For PrP and GFAP staining Mouse on Mouse Elite Peroxidase Kit (Vector Laboratories) was used according to the manufacturer’s instructions. For Iba1 staining 2,5% donkey normal serum (S30, Merck Millipore) was used for blocking (30min), followed by incubation with the primary antibody (30-60min), biotinylated secondary antibody (30min) and AB- Complex (30 min). Staining of the epitope was done with DAB with Nickel for all stainings (DAB Peroxidase (HRP) Substrate Kit with Nickel, Vector Laboratories; 5-10 min). A counterstain with hematoxylin (H-3404, Vector Laboratories) was performed (10 s) and sections were dehydrated with graded dilutions of ethanol and xylene (each 5 min). For each time point, all brain sections were treated at the same time with the same solutions and materials. Pictures of sections were made with an AxioCam camera mounted onto a Zeiss AXIO Observer.A1 microscope with Zen 2012 software used with the same microscope imaging parameters for related stainings and magnifications. Primary antibodies: Anti-GFAP Clone GA5 (MAB360), 1:5000, Merck Millipore; Prion Protein Monoclonal Antibody SAF84, 1:200, Cayman Chemical; Rabbit Anti Iba1 for ICC, 1:200, Waco; Rabbit anti Rps21, 1:100, Bethyl (A305-070A); secondary antibodies: Biotin-SP-conjugated AffiniPure Donkey Anti-Rabbit IgG, 1:500, Jackson Immuno Research.

### Immunofluorescence

In order to assess the cell type-specificity of RiboTag experiments, we processed additional mice for immunohistological evaluation. Mice were perfused transcardially with saline followed by 10% neutral buffered formalin solution (Sigma) and immersion-postfixed in the same solution overnight at 4C. After cryoprotection in 30% sucrose in PBS, 40 µm coronal cryosections were taken and sections stored until use in 50% 0.1 M PBS, 30% ethylene glycol, and 20% glycerol at -20C. Sections at the level of AP-1.46 (Paxinos and Franklin, 2nd edition) were selected, and double incubated overnight at 4°C in primary antibodies against HA (1:200, 3F10, Sigma) and the respective marker antibody (Somatostatin, 1:200, HPA019472, Atlas; Sox9, 1:500, AB5535, Millipore; NeuN, 1:500, MAB377, Millipore; Parvalbumin, 1:200, Novus, NB-120-11427; Satb2, 1:200, Abcam, 92446; GFAP, 1:500, Dako, Z0334; Iba1, 1:100, Wako, 019-19741; Gad67, 1:2000, Acris, AP15571PU). The double incubation for HA and Gad67 was preceded by a 45 min antigene retrieval in pH6 citrate buffer using a vegetable steamer (15 min + 30 min cooling to RT), quenching of peroxidases in 0.3% H2O2, and avidin-biotin block (SP-2001, Vector). Visualization occurred by double incubation in fluorescent secondary antibodies from Jackson Immun Research against rat IgG (712-165-153 or 712-545-153) and appropriate second antibody (mouse, 715-545-151; rabbit, 711-225-152 or 711-585-152). For Gad67 labeling we instead used a biotinylated secondary antibody (711-065-152) followed by tyramide signal amplification (T20935, Thermo Fisher).

## References

Ansoleaga, B., Garcia-Esparcia, P., Llorens, F., Hernandez-Ortega, K., Carmona Tech, M., Antonio Del Rio, J., . . . Ferrer, I. (2016). Altered Mitochondria, Protein Synthesis Machinery, and Purine Metabolism Are Molecular Contributors to the Pathogenesis of Creutzfeldt-Jakob Disease. J Neuropathol Exp Neurol, 75(8), 755–769. doi:10.1093/jnen/nlw048

Battle, A., Khan, Z., Wang, S. H., Mitrano, A., Ford, M. J., Pritchard, J. K., & Gilad, Y. (2015). Genomic variation. Impact of regulatory variation from RNA to protein. Science, 347(6222), 664–667. doi:10.1126/science.1260793

Boisvert, M. M., Erikson, G. A., Shokhirev, M. N., & Allen, N. J. (2018). The Aging Astrocyte Transcriptome from Multiple Regions of the Mouse Brain. Cell Rep, 22(1), 269–285. doi:10.1016/j.celrep.2017.12.039

Carroll, T., Guha, S., Nehrke, K., & Johnson, G. V. W. (2021). Tau Post-Translational Modifications: Potentiators of Selective Vulnerability in Sporadic Alzheimer’s Disease. Biology (Basel), 10(10). doi:10.3390/biology10101047

Carter, B., Justin, H. S., Gulick, D., & Gamsby, J. J. (2021). The Molecular Clock and Neurodegenerative Disease: A Stressful Time. Front Mol Biosci, 8, 644747. doi:10.3389/fmolb.2021.644747

Chen, T., Zhu, J., Wang, Y. H., & Hang, C. H. (2020). Arc silence aggravates traumatic neuronal injury via mGluR1-mediated ER stress and necroptosis. Cell Death Dis, 11(1), 4. doi:10.1038/s41419-019-2198-5

Corbett, G. T., Wang, Z., Hong, W., Colom-Cadena, M., Rose, J., Liao, M., . . . Walsh, D.M. (2020). PrP is a central player in toxicity mediated by soluble aggregates of neurodegeneration-causing proteins. Acta Neuropathol, 139(3), 503–526. doi:10.1007/s00401-019-02114-9

Delarue, M., Brittingham, G. P., Pfeffer, S., Surovtsev, I. V., Pinglay, S., Kennedy, K. J., . . . Holt, L. J. (2018). mTORC1 Controls Phase Separation and the Biophysical Properties of the Cytoplasm by Tuning Crowding. Cell, 174(2), 338–349 e320. doi:10.1016/j.cell.2018.05.042

Dittrich, L., Morairty, S. R., Warrier, D. R., & Kilduff, T. S. (2015). Homeostatic sleep pressure is the primary factor for activation of cortical nNOS/NK1 neurons. Neuropsychopharmacology, 40(3), 632–639. doi:10.1038/npp.2014.212

Dittrich, L., Petese, A., & Jackson, W. S. (2017). The natural Disc1-deletion present in several inbred mouse strains does not affect sleep. Sci Rep, 7(1), 5665. doi:10.1038/s41598-017-06015-3

Edgar, R., Domrachev, M., & Lash, A. E. (2002). Gene Expression Omnibus: NCBI gene expression and hybridization array data repository. Nucleic Acids Res, 30(1), 207–210. doi:10.1093/nar/30.1.207

Evers, T. M. J., Holt, L. J., Alberti, S., & Mashaghi, A. (2021). Reciprocal regulation of cellular mechanics and metabolism. Nat Metab, 3(4), 456–468. doi:10.1038/s42255-021-00384-w

Ferrer, I., Casas, R., & Rivera, R. (1993). Parvalbumin-immunoreactive cortical neurons in Creutzfeldt-Jakob disease. Ann Neurol, 34(6), 864–866. doi:10.1002/ana.410340617

Fifel, K., & Videnovic, A. (2020). Circadian alterations in patients with neurodegenerative diseases: Neuropathological basis of underlying network mechanisms. Neurobiol Dis, 144, 105029. doi:10.1016/j.nbd.2020.105029

Fornasiero, E. F., Mandad, S., Wildhagen, H., Alevra, M., Rammner, B., Keihani, S., .. . Rizzoli, S.O. (2018). Precisely measured protein lifetimes in the mouse brain reveal differences across tissues and subcellular fractions. Nat Commun, 9(1), 4230. doi:10.1038/s41467-018-06519-0

Franko, E., Wehner, T., Joly, O., Lowe, J., Porter, M. C., Kenny, J., . . . Mead, S. (2016). Quantitative EEG parameters correlate with the progression of human prion diseases. J Neurol Neurosurg Psychiatry, 87(10), 1061–1067. doi:10.1136/jnnp-2016-313501

Frau-Mendez, M. A., Fernandez-Vega, I., Ansoleaga, B., Blanco Tech, R., Carmona Tech, M., Antonio Del Rio, J., . . . Ferrer, I. (2017). Fatal familial insomnia: mitochondrial and protein synthesis machinery decline in the mediodorsal thalamus. Brain Pathol, 27(1), 95–106. doi:10.1111/bpa.12408

Fu, H., Hardy, J., & Duff, K. E. (2018). Selective vulnerability in neurodegenerative diseases. Nat Neurosci, 21(10), 1350–1358. doi:10.1038/s41593-018-0221-2

Gawinecka, J., Nowak, M., Carimalo, J., Cardone, F., Asif, A. R., Wemheuer, W. M., .. . Zerr, I. (2013). Subtype-specific synaptic proteome alterations in sporadic Creutzfeldt-Jakob disease. J Alzheimers Dis, 37(1), 51–61. doi:10.3233/JAD-130455

Ghiassian, S. D., Menche, J., & Barabasi, A. L. (2015). A DIseAse MOdule Detection (DIAMOnD) algorithm derived from a systematic analysis of connectivity patterns of disease proteins in the human interactome. PLoS Comput Biol, 11(4), e1004120. doi:10.1371/journal.pcbi.1004120

Gibbings, D., Leblanc, P., Jay, F., Pontier, D., Michel, F., Schwab, Y., . . . Voinnet, O. (2012). Human prion protein binds Argonaute and promotes accumulation of microRNA effector complexes. Nat Struct Mol Biol, 19(5), 517–524, S511. doi:10.1038/nsmb.2273

Goedert, M., Clavaguera, F., & Tolnay, M. (2010). The propagation of prion-like protein inclusions in neurodegenerative diseases. Trends Neurosci, 33(7), 317–325. doi:10.1016/j.tins.2010.04.003

Guentchev, M., Groschup, M. H., Kordek, R., Liberski, P. P., & Budka, H. (1998). Severe, early and selective loss of a subpopulation of GABAergic inhibitory neurons in experimental transmissible spongiform encephalopathies. Brain Pathol, 8(4), 615–623.

Guentchev, M., Hainfellner, J. A., Trabattoni, G. R., & Budka, H. (1997). Distribution of parvalbumin-immunoreactive neurons in brain correlates with hippocampal and temporal cortical pathology in Creutzfeldt-Jakob disease. J Neuropathol Exp Neurol, 56(10), 1119–1124.

Guentchev, M., Wanschitz, J., Voigtlander, T., Flicker, H., & Budka, H. (1999). Selective neuronal vulnerability in human prion diseases. Fatal familial insomnia differs from other types of prion diseases. Am J Pathol, 155(5), 1453–1457. doi:10.1016/S0002-9440(10)65459-4

Halder, R., Hennion, M., Vidal, R. O., Shomroni, O., Rahman, R. U., Rajput, A., . . . Bonn, S. (2016). DNA methylation changes in plasticity genes accompany the formation and maintenance of memory. Nat Neurosci, 19(1), 102–110. doi:10.1038/nn.4194

Hetz, C., Zhang, K., & Kaufman, R. J. (2020). Mechanisms, regulation and functions of the unfolded protein response. Nat Rev Mol Cell Biol, 21(8), 421–438. doi:10.1038/s41580-020-0250-z

Hippenmeyer, S., Vrieseling, E., Sigrist, M., Portmann, T., Laengle, C., Ladle, D. R., & Arber, S. (2005). A developmental switch in the response of DRG neurons to ETS transcription factor signaling. PLoS Biol, 3(5), e159. doi:10.1371/journal.pbio.0030159

Hol, E. M., Roelofs, R. F., Moraal, E., Sonnemans, M. a. F., Sluijs, J. A., Proper, E. A.,. . . van Leeuwen, F. W. (2003). Neuronal expression of GFAP in patients with Alzheimer pathology and identification of novel GFAP splice forms. Molecular Psychiatry, 8(9), 786–796. doi:10.1038/sj.mp.4001379

Hoshino, A., Helwig, M., Rezaei, S., Berridge, C., Eriksen, J. L., & Lindberg, I. (2014). A novel function for proSAAS as an amyloid anti-aggregant in Alzheimer’s disease. J Neurochem, 128(3), 419–430. doi:10.1111/jnc.12454

Iliff, J. J., Wang, M., Liao, Y., Plogg, B. A., Peng, W., Gundersen, G. A., . . . Nedergaard, M. (2012). A paravascular pathway facilitates CSF flow through the brain parenchyma and the clearance of interstitial solutes, including amyloid beta. Sci Transl Med, 4(147), 147ra111. doi:10.1126/scitranslmed.3003748

Jackson, W. S. (2014). Selective vulnerability to neurodegenerative disease: the curious case of Prion Protein. Dis Model Mech, 7(1), 21–29. doi:10.1242/dmm.012146

Jackson, W. S., Borkowski, A. W., Faas, H., Steele, A. D., King, O. D., Watson, N., . .. Lindquist, S. (2009). Spontaneous generation of prion infectivity in fatal familial insomnia knockin mice. Neuron, 63(4), 438–450. doi:S0896-6273(09)00581-9 [pii]10.1016/j.neuron.2009.07.026

Jackson, W. S., Borkowski, A. W., Watson, N. E., King, O. D., Faas, H., Jasanoff, A., & Lindquist, S. (2013). Profoundly different prion diseases in knock-in mice carrying single PrP codon substitutions associated with human diseases. Proc Natl Acad Sci U S A, 110(36), 14759–14764. doi:10.1073/pnas.1312006110

Ji, L., Chauhan, A., Wegiel, J., Essa, M. M., & Chauhan, V. (2009). Gelsolin is proteolytically cleaved in the brains of individuals with Alzheimer’s disease. J Alzheimers Dis, 18(1), 105–111. doi:10.3233/JAD-2009-1127

Kaczmarczyk, L., Bansal, V., Rajput, A., Rahman, R. U., Krzyzak, W., Degen, J., . . . Jackson, W. S. (2019). Tagger-A Swiss army knife for multiomics to dissect cell type-specific mechanisms of gene expression in mice. PLoS Biol, 17(8), e3000374. doi:10.1371/journal.pbio.3000374

Kaczmarczyk, L., Reichenbach, N., Blank, N., Jonson, M., Dittrich, L., Petzold, G. C., & Jackson, W. S. (2021). Slc1a3-2A-CreERT2 mice reveal unique features of Bergmann glia and augment a growing collection of Cre drivers and effectors in the 129S4 genetic background Sci Rep, 11(1), 5412. doi:10.1038/s41598-021-84887-2

Kang, S. S., Ebbert, M. T. W., Baker, K. E., Cook, C., Wang, X., Sens, J. P., . . . Fryer, J. D. (2018). Microglial translational profiling reveals a convergent APOE pathway from aging, amyloid, and tau. J Exp Med, 215(9), 2235–2245. doi:10.1084/jem.20180653

Kretz, M., Euwens, C., Hombach, S., Eckardt, D., Teubner, B., Traub, O., . . . Ott, T. (2003). Altered connexin expression and wound healing in the epidermis of connexin-deficient mice. J Cell Sci, 116(Pt 16), 3443–3452. doi:10.1242/jcs.00638

Langmead, B., & Salzberg, S. L. (2012). Fast gapped-read alignment with Bowtie 2. Nat Methods, 9(4), 357–359. doi:10.1038/nmeth.1923

Lau, A., So, R. W. L., Lau, H. H. C., Sang, J. C., Ruiz-Riquelme, A., Fleck, S. C., . . . Watts, J. C. (2020). alpha-Synuclein strains target distinct brain regions and cell types. Nat Neurosci, 23(1), 21–31. doi:10.1038/s41593-019-0541-x

Leng, Y., Musiek, E. S., Hu, K., Cappuccio, F. P., & Yaffe, K. (2019). Association between circadian rhythms and neurodegenerative diseases. Lancet Neurol, 18(3), 307–318. doi:10.1016/S1474-4422(18)30461-7

Leuchter, M. K., Donzis, E. J., Cepeda, C., Hunter, A. M., Estrada-Sanchez, A. M., Cook, I. A., . . . Leuchter, A. F. (2017). Quantitative Electroencephalographic Biomarkers in Preclinical and Human Studies of Huntington’s Disease: Are They Fit-for-Purpose for Treatment Development? Front Neurol, 8, 91. doi:10.3389/fneur.2017.00091

Li, B., & Dewey, C. N. (2011). RSEM: accurate transcript quantification from RNA-Seq data with or without a reference genome. BMC Bioinformatics, 12, 323. doi:10.1186/1471-2105-12-323

Liao, Y., Wang, J., Jaehnig, E. J., Shi, Z., & Zhang, B. (2019). WebGestalt 2019: gene set analysis toolkit with revamped UIs and APIs. Nucleic Acids Res, 47(W1), W199–W205. doi:10.1093/nar/gkz401

Liberzon, A., Subramanian, A., Pinchback, R., Thorvaldsdottir, H., Tamayo, P., & Mesirov, J. P. (2011). Molecular signatures database (MSigDB) 3.0. Bioinformatics, 27(12), 1739–1740. doi:10.1093/bioinformatics/btr260

Love, M. I., Huber, W., & Anders, S. (2014). Moderated estimation of fold change and dispersion for RNA-seq data with DESeq2. Genome Biol, 15(12), 550. doi:10.1186/s13059-014-0550-8

Magger, O., Waldman, Y. Y., Ruppin, E., & Sharan, R. (2012). Enhancing the prioritization of disease-causing genes through tissue specific protein interaction networks. PLoS Comput Biol, 8(9), e1002690. doi:10.1371/journal.pcbi.1002690

Majer, A., Medina, S. J., Niu, Y., Abrenica, B., Manguiat, K. J., Frost, K. L., . . . Booth, S. A. (2012). Early mechanisms of pathobiology are revealed by transcriptional temporal dynamics in hippocampal CA1 neurons of prion infected mice. PLoS Pathog, 8(11), e1003002. doi:10.1371/journal.ppat.1003002

Majer, A., Medina, S. J., Sorensen, D., Martin, M. J., Frost, K. L., Phillipson, C., . . . Booth, S. A. (2019). The cell type resolved mouse transcriptome in neuron-enriched brain tissues from the hippocampus and cerebellum during prion disease. Sci Rep, 9(1), 1099. doi:10.1038/s41598-018-37715-z

Makarava, N., Chang, J. C., Kushwaha, R., & Baskakov, I. V. (2019). Region-Specific Response of Astrocytes to Prion Infection. Front Neurosci, 13, 1048. doi:10.3389/fnins.2019.01048

Makarava, N., Mychko, O., Chang, J. C., Molesworth, K., & Baskakov, I. V. (2021). The degree of astrocyte activation is predictive of the incubation time to prion disease. Acta Neuropathol Commun, 9(1), 87. doi:10.1186/s40478-021-01192-9

Mathys, H., Davila-Velderrain, J., Peng, Z., Gao, F., Mohammadi, S., Young, J. Z., . . . Tsai, L. H. (2019). Single-cell transcriptomic analysis of Alzheimer’s disease. Nature, 570(7761), 332–337. doi:10.1038/s41586-019-1195-2

Mattsson, N., Schott, J. M., Hardy, J., Turner, M. R., & Zetterberg, H. (2016). Selective vulnerability in neurodegeneration: insights from clinical variants of Alzheimer’s disease. J Neurol Neurosurg Psychiatry, 87(9), 1000–1004. doi:10.1136/jnnp-2015-311321

Meghdadi, A. H., Stevanovic Karic, M., McConnell, M., Rupp, G., Richard, C., Hamilton, J., . . . Berka, C. (2021). Resting state EEG biomarkers of cognitive decline associated with Alzheimer’s disease and mild cognitive impairment. PLoS One, 16(2), e0244180. doi:10.1371/journal.pone.0244180

Millard, S. M., Heng, O., Opperman, K. S., Sehgal, A., Irvine, K. M., Kaur, S., . . . Pettit, A. R. (2021). Fragmentation of tissue-resident macrophages during isolation confounds analysis of single-cell preparations from mouse hematopoietic tissues. Cell Rep, 37(8), 110058. doi:10.1016/j.celrep.2021.110058

Minatohara, K., Akiyoshi, M., & Okuno, H. (2015). Role of Immediate-Early Genes in Synaptic Plasticity and Neuronal Ensembles Underlying the Memory Trace. Front Mol Neurosci, 8, 78. doi:10.3389/fnmol.2015.00078

Mong, J. A., Baker, F. C., Mahoney, M. M., Paul, K. N., Schwartz, M. D., Semba, K., & Silver, R. (2011). Sleep, rhythms, and the endocrine brain: influence of sex and gonadal hormones. J Neurosci, 31(45), 16107–16116. doi:10.1523/JNEUROSCI.4175-11.2011

Mong, J. A., & Cusmano, D. M. (2016). Sex differences in sleep: impact of biological sex and sex steroids. Philos Trans R Soc Lond B Biol Sci, 371(1688), 20150110. doi:10.1098/rstb.2015.0110

Morairty, S. R., Dittrich, L., Pasumarthi, R. K., Valladao, D., Heiss, J. E., Gerashchenko, D., & Kilduff, T. S. (2013). A role for cortical nNOS/NK1 neurons in coupling homeostatic sleep drive to EEG slow wave activity. Proc Natl Acad Sci U S A, 110(50), 20272–20277. doi:10.1073/pnas.1314762110

Moreno, J. A., Radford, H., Peretti, D., Steinert, J. R., Verity, N., Martin, M. G., . . . Mallucci, G. R. (2012). Sustained translational repression by eIF2alpha-P mediates prion neurodegeneration. Nature, 485(7399), 507–511. doi:10.1038/nature11058

Musiek, E. S., & Holtzman, D. M. (2016). Mechanisms linking circadian clocks, sleep, and neurodegeneration. Science, 354(6315), 1004–1008. doi:10.1126/science.aah4968

Musiek, E. S., Lim, M. M., Yang, G., Bauer, A. Q., Qi, L., Lee, Y., . . . Fitzgerald, G. A. (2013). Circadian clock proteins regulate neuronal redox homeostasis and neurodegeneration. J Clin Invest, 123(12), 5389–5400. doi:10.1172/JCI70317

Parks, G. S., Warrier, D. R., Dittrich, L., Schwartz, M. D., Palmerston, J. B., Neylan, T. C., . . . Kilduff, T. S. (2016). The Dual Hypocretin Receptor Antagonist Almorexant is Permissive for Activation of Wake-Promoting Systems. Neuropsychopharmacology, 41(4), 1144–1155. doi:10.1038/npp.2015.256

Pfrieger, F. W., & Slezak, M. (2012). Genetic approaches to study glial cells in the rodent brain. Glia, 60(5), 681–701. doi:10.1002/glia.22283

Preiss, T. (2016). All Ribosomes Are Created Equal. Really? Trends Biochem Sci, 41(2), 121–123. doi:10.1016/j.tibs.2015.11.009

Sanz, E., Yang, L., Su, T., Morris, D. R., McKnight, G. S., & Amieux, P. S. (2009). Cell-type-specific isolation of ribosome-associated mRNA from complex tissues. Proc Natl Acad Sci U S A, 106(33), 13939–13944. doi:0907143106 [pii] 10.1073/pnas.0907143106

Scheckel, C., Imeri, M., Schwarz, P., & Aguzzi, A. (2020). Ribosomal profiling during prion disease uncovers progressive translational derangement in glia but not in neurons. Elife, 9. doi:10.7554/eLife.62911

Shi, Z., Fujii, K., Kovary, K. M., Genuth, N. R., Rost, H. L., Teruel, M. N., & Barna, M. (2017). Heterogeneous Ribosomes Preferentially Translate Distinct Subpools of mRNAs Genome-wide. Mol Cell, 67(1), 71–83 e77. doi:10.1016/j.molcel.2017.05.021

Sinha, A., Kushwaha, R., Molesworth, K., Mychko, O., Makarava, N., & Baskakov, I. V. (2021). Phagocytic Activities of Reactive Microglia and Astrocytes Associated with Prion Diseases Are Dysregulated in Opposite Directions. Cells, 10(7). doi:10.3390/cells10071728

Smith, H. L., Freeman, O. J., Butcher, A. J., Holmqvist, S., Humoud, I., Schatzl, T., . . . Mallucci, G. R. (2020). Astrocyte Unfolded Protein Response Induces a Specific Reactivity State that Causes Non-Cell-Autonomous Neuronal Degeneration. Neuron, 105(5), 855–866 e855. doi:10.1016/j.neuron.2019.12.014

Sofroniew, M. V., & Vinters, H. V. (2010). Astrocytes: biology and pathology. Acta Neuropathol, 119(1), 7–35. doi:10.1007/s00401-009-0619-8

Sorce, S., Nuvolone, M., Russo, G., Chincisan, A., Heinzer, D., Avar, M., . . . Aguzzi, A. (2020). Genome-wide transcriptomics identifies an early preclinical signature of prion infection. PLoS Pathog, 16(6), e1008653. doi:10.1371/journal.ppat.1008653

Steward, O., Wallace, C. S., Lyford, G. L., & Worley, P. F. (1998). Synaptic activation causes the mRNA for the IEG Arc to localize selectively near activated postsynaptic sites on dendrites. Neuron, 21(4), 741–751. doi:10.1016/s0896-6273(00)80591-7

Sun, S., Sun, Y., Ling, S. C., Ferraiuolo, L., McAlonis-Downes, M., Zou, Y., . . . Cleveland, D. W. (2015). Translational profiling identifies a cascade of damage initiated in motor neurons and spreading to glia in mutant SOD1-mediated ALS. Proc Natl Acad Sci U S A, 112(50), E6993–7002. doi:10.1073/pnas.1520639112

Szklarczyk, D., Gable, A. L., Nastou, K. C., Lyon, D., Kirsch, R., Pyysalo, S., . . . von Mering, C. (2021). The STRING database in 2021: customizable protein-protein networks, and functional characterization of user-uploaded gene/measurement sets. Nucleic Acids Res, 49(D1), D605–D612. doi:10.1093/nar/gkaa1074

Taniguchi, H. (2014). Genetic dissection of GABAergic neural circuits in mouse neocortex. Front Cell Neurosci, 8, 8. doi:10.3389/fncel.2014.00008

Taniguchi, H., He, M., Wu, P., Kim, S., Paik, R., Sugino, K., . . . Huang, Z. J. (2011). A resource of Cre driver lines for genetic targeting of GABAergic neurons in cerebral cortex. Neuron, 71(6), 995–1013. doi:10.1016/j.neuron.2011.07.026

Tebbenkamp, A. T., & Borchelt, D. R. (2010). Analysis of chaperone mRNA expression in the adult mouse brain by meta analysis of the Allen Brain Atlas. PLoS One, 5(10), e13675. doi:10.1371/journal.pone.0013675

Vanni, S., Moda, F., Zattoni, M., Bistaffa, E., De Cecco, E., Rossi, M., . . . Legname, G. (2017). Differential overexpression of SERPINA3 in human prion diseases. Sci Rep, 7(1), 15637. doi:10.1038/s41598-017-15778-8

Vincenti, J. E., Murphy, L., Grabert, K., McColl, B. W., Cancellotti, E., Freeman, T. C., & Manson, J. C. (2015). Defining the Microglia Response during the Time Course of Chronic Neurodegeneration. J Virol, 90(6), 3003–3017. doi:10.1128/JVI.02613-15

Vong, L., Ye, C., Yang, Z., Choi, B., Chua, S., Jr., & Lowell, B. B. (2011). Leptin action on GABAergic neurons prevents obesity and reduces inhibitory tone to POMC neurons. Neuron, 71(1), 142–154. doi:10.1016/j.neuron.2011.05.028

Walker, L. C., Schelle, J., & Jucker, M. (2016). The Prion-Like Properties of Amyloid-beta Assemblies: Implications for Alzheimer’s Disease. Cold Spring Harb Perspect Med, 6(7). doi:10.1101/cshperspect.a024398

Watts, J. C., & Prusiner, S. B. (2014). Mouse Models for Studying the Formation and Propagation of Prions. J Biol Chem. doi:10.1074/jbc.R114.550707

Xue, S., & Barna, M. (2012). Specialized ribosomes: a new frontier in gene regulation and organismal biology. Nat Rev Mol Cell Biol, 13(6), 355–369. doi:10.1038/nrm3359

Yan, X. X., Jeromin, A., & Jeromin, A. (2012). Spectrin Breakdown Products (SBDPs) as Potential Biomarkers for Neurodegenerative Diseases. Curr Transl Geriatr Exp Gerontol Rep, 1(2), 85–93. doi:10.1007/s13670-012-0009-2

Yu, G., Wang, L. G., Han, Y., & He, Q. Y. (2012). clusterProfiler: an R package for comparing biological themes among gene clusters. OMICS, 16(5), 284–287. doi:10.1089/omi.2011.0118

Zhang, S., Zhao, J., Zhang, Y., Zhang, Y., Cai, F., Wang, L., & Song, W. (2019). Upregulation of MIF as a defense mechanism and a biomarker of Alzheimer’s disease. Alzheimers Res Ther, 11(1), 54. doi:10.1186/s13195-019-0508-x

Zhu, C., Herrmann, U. S., Falsig, J., Abakumova, I., Nuvolone, M., Schwarz, P., . . . Aguzzi, A. (2016). A neuroprotective role for microglia in prion diseases. J Exp Med, 213(6), 1047–1059. doi:10.1084/jem.20151000

